# Neutralizing monoclonal antibodies elicited by mosaic RBD nanoparticles bind conserved sarbecovirus epitopes

**DOI:** 10.1101/2022.06.28.497989

**Authors:** Chengcheng Fan, Alexander A. Cohen, Miso Park, Alfur Fu-Hsin Hung, Jennifer R. Keeffe, Priyanthi N.P. Gnanapragasam, Yu E. Lee, Leesa M. Kakutani, Ziyan Wu, Kathryn E. Malecek, John C. Williams, Pamela J. Bjorkman

## Abstract

Protection from SARS-related coronaviruses with spillover potential and SARS-CoV-2 variants could prevent and/or end pandemics. We show that mice immunized with nanoparticles co-displaying spike receptor-binding domains (RBDs) from eight sarbecoviruses (mosaic-8 RBD-nanoparticles) efficiently elicit cross-reactive anti-sarbecovirus antibodies against conserved class 1/4 and class 3 RBD epitopes. Monoclonal antibodies (mAbs) identified from initial screening of <10,000 single B-cells secreting IgGs binding two or more sarbecovirus RBDs showed cross-reactive binding and neutralization of SARS-CoV-2 variants and animal sarbecoviruses. Single-particle cryo-EM structures of antibody–spike complexes, including a Fab-Omicron complex, mapped neutralizing mAbs to conserved class 1/4 RBD epitopes and revealed neutralization mechanisms, potentials for intra-spike trimer crosslinking by single IgGs, and induced changes in trimer upon Fab binding. In addition, we identified a mAb resembling Bebtelovimab, an EUA-approved human class 3 anti-RBD mAb. These results support using mosaic RBD-nanoparticles to identify therapeutic pan-sarbecovirus and pan-variant mAbs and to elicit them by vaccination.

## Introduction

Spillover of animal SARS-like betacoronaviruses (sarbecoviruses) resulted in two human health emergencies in the past 20 years: the SARS-CoV epidemic in the early 2000s and the current COVID-19 pandemic caused by SARS-CoV-2. Large coronavirus reservoirs in bats are predictive of future cross-species transmission (Menachery et al., 2015; Menachery et al., 2016; Zhou et al., 2021), necessitating a vaccine that could protect against emerging coronaviruses. In addition, SARS-CoV-2 variants have been discovered throughout the current pandemic, with the Alpha, Beta, Delta, Gamma, and Omicron lineages designated as variants of concern (VOCs) due to apparent increased transmissibility and/or resistance to neutralizing antibodies elicited by infection or vaccination (Burki, 2021; Liu et al., 2021; Planas et al., 2021; Washington et al., 2021). In the case of Omicron VOCs, a large number of substitutions in the SARS-CoV-2 spike protein receptor-binding domain (RBD), the major target of neutralizing antibodies (Barnes et al., 2020a; Brouwer et al., 2020; Cao et al., 2020; Kreer et al., 2020; Liu et al., 2020b; Piccoli et al., 2020; Pinto et al., 2020; Robbiani et al., 2020; Rogers et al., 2020; Seydoux et al., 2020; Zost et al., 2020) and detectable cross-variant neutralization (Bowen et al., 2021), results in reduced efficacies of vaccines and therapeutic monoclonal antibodies (mAbs) (Liu et al., 2021; Starr et al., 2021).

Comparison of the variability of RBDs across sarbecoviruses (Figure S1) and within SARS-CoV-2 VOCs and variants of interest (VOIs) suggest that vaccines and mAbs targeting the more conserved neutralizing antibody epitopes (class 4 and class 1/4; nomenclature from (Barnes et al., 2020a; Jette et al., 2021)) (Figure 1A,B) could protect against present and future SARS-CoV-2 VOCs and prevent future sarbecovirus spillover events from causing another epidemic or pandemic. By contrast, antibodies targeting the less conserved class 1 and class 2 RBD epitopes that directly overlap with the binding footprint for human ACE2, the SARS-CoV-2 host receptor, recognize a portion of the RBD that exhibits sequence variability between sarbecoviruses (Barnes et al., 2020a) (Figure 1A), which is also where VOC and VOI substitutions accumulate (Figure 1B; Figure S2). Class 3 RBD epitopes are more conserved than class 1 and class 2 epitopes but exhibit some variation across sarbecoviruses (Figure 1B; Figure S2), suggesting the potential for continued variability amongst SARS-CoV-2 VOCs.

**Figure 1.**
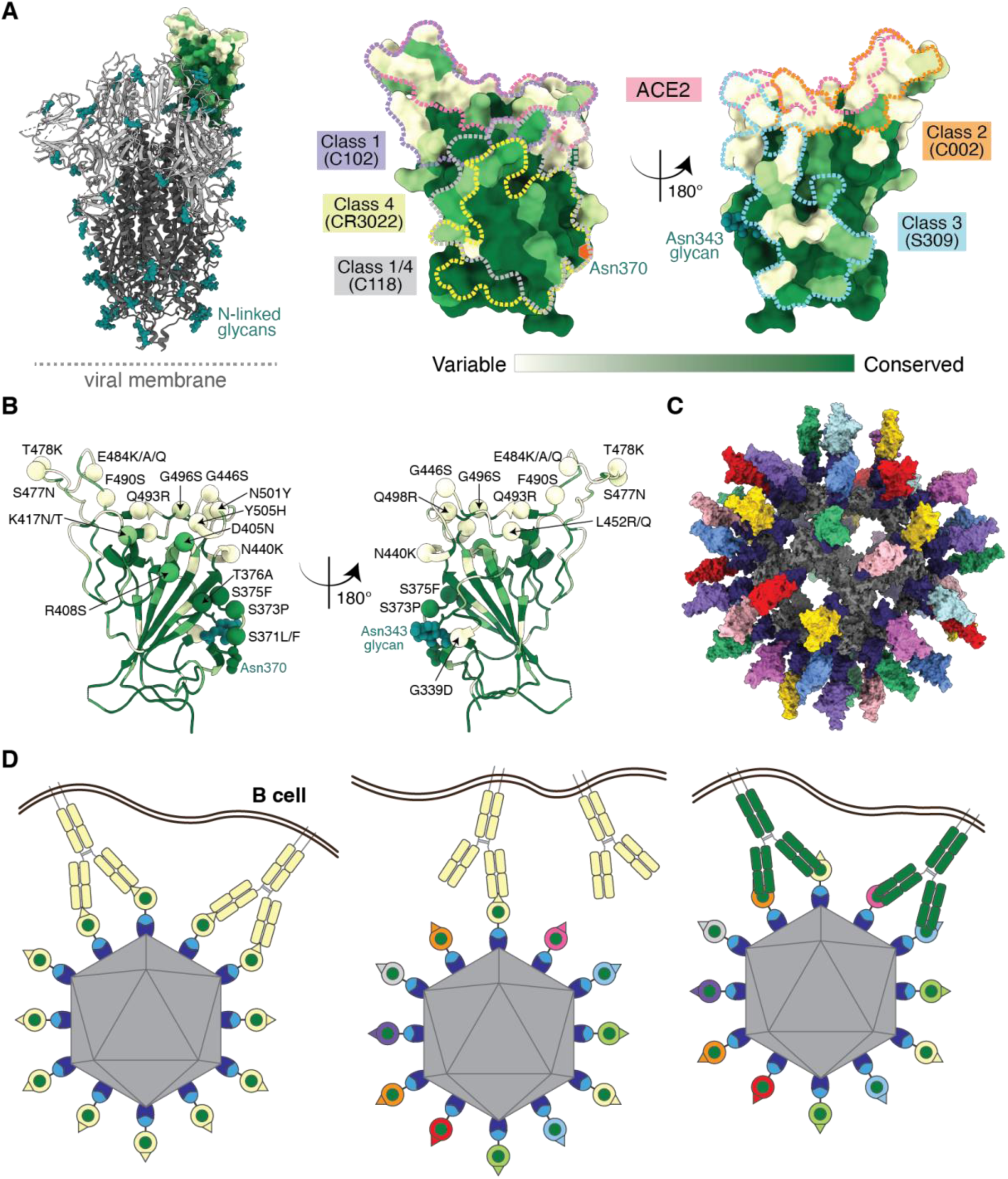
Utilizing antibody avidity effects suggests a strategy to target antibodies to conserved regions of sarbecovirus RBDs. (**A**) Left: Structure of SARS-CoV-2 spike trimer (PDB 6VYB) with one RBD in an “up” position. Right: Sequence conservation of 16 sarbecovirus RBDs (Figure S1) calculated by the ConSurf Database (Landau et al., 2005) plotted on a surface representation of the RBD structure (PDB 7BZ5). Class 1, 2, 3, 4, and 1/4 epitopes are outlined in different colored dots using information from structures of the representative monoclonal antibodies bound to RBD or spike trimer (C102: PDB 7K8M; C002: PDB 7K8T, S309: PDB 7JX3; CR3022: PDB 7LOP; C118: PDB 7RKV). (**B**) RBD mutations of 13 SARS-CoV-2 VOCs and VOIs (https://viralzone.expasy.org/9556) plotted onto the RBD structure (PDB 7BZ5) as spheres that are colored according to the variability gradient in panel A. The N-linked glycan at position 343 of SARS-CoV-2 RBD is shown as teal spheres, and a potential N-linked glycosylation site at position 370 (SARS-CoV-2 numbering) found in some sarbecovirus RBDs but not in the SARS-CoV-2 RBD is indicated by an orange hexagon. (**C**) Structural model of mosaic-8 nanoparticle formed by SpyCatcher-mi3 and eight SpyTagged RBDs made using coordinates of an RBD (PDB 7SC1), mi3 (PDB 7B3Y), and SpyCatcher (PDB 4MLI). (**D**) Hypothesis for preferential stimulation of B cells with that encode cross-reactive antibodies by mosaic (right) versus homotypic (left) RBD nanoparticles. Left: Yellow B cell receptors recognizing an accessible strain-specific epitope (yellow triangle) can crosslink between adjacent RBDs on a homotypic nanoparticle to enhance binding through avidity effects. Middle: Yellow B cell receptors against a strain-specific orange epitope cannot crosslink between adjacent RBDs on a mosaic RBD nanoparticle that presents different versions of the epitope (colored triangles). Right: Green cross-reactive B cell receptors can crosslink between a conserved epitope (green circles) on adjacent RBDs in a mosaic RBD nanoparticle to enhance binding to a more occluded, but conserved, epitope through avidity effects.

We previously described a vaccine approach involving simultaneous display of sarbecovirus RBDs on protein-based nanoparticles (Figure 1C) that showed enhanced heterologous binding, neutralization, and protection from sarbecovirus challenges compared with homotypic (SARS-CoV-2 RBD–only) nanoparticles in animal models (Cohen et al., 2021; Cohen et al., 2022). The hypothesis behind enhanced elicitation of cross-reactive antibodies by mosaic RBD-nanoparticles is that B cell receptors (BCRs) that recognize conserved, but not readily accessible, RBD epitopes would be stimulated to proliferate to produce cross-reactive Abs through bivalent binding of BCRs to adjacent RBDs, whereas BCRs that bind to strain-specific epitopes could not bind bivalently to adjacent RBDs since these epitopes differ when RBDs are arranged randomly on a nanoparticle (Figure 1D) (Cohen et al., 2022). By contrast, homotypic RBD-nanoparticles are predicted to stimulate BCRs against readily accessible, immunodominant strain-specific epitopes presented on all RBDs. Of relevance to the mAb potentials for sarbecovirus cross-reactivity, the more conserved class 4 and class 1/4 epitopes targeted by polyclonal antibodies in mosaic-8 RBD-nanoparticle antisera are unlikely to vary in SARS-CoV-2 VOCs because they contact other portions of the spike trimer, by contrast to the class 1 and 2 RBD epitope regions targeted by homotypic SARS-CoV-2 RBD-nanoparticle antisera, which (in common with the class 3 RBD region) are not involved in contacts with non-RBD portions of spike (Cohen et al., 2022). Although the epitopes of both class 4 and class 1/4 anti-RBD mAbs fall outside of the ACE2-binding footprint within class 1 and class 2 RBD epitopes, these mAbs can directly compete with ACE2 for binding through steric effects, thereby rationalizing their increased neutralization potencies (Jette et al., 2021; Liu et al., 2020a; Tortorici et al., 2021).

Here we investigated the RBD epitopes of mAbs isolated from mosaic RBD- and homotypic RBD-immunized mice to characterize the antibody response to RBD nanoparticles. Binding and neutralization results, together with cryo-EM structures of antibody Fab-spike trimer complexes, suggested that the mosaic RBD-nanoparticle vaccine approach works as designed to target conserved epitopes, and could be used both for more broadly protective vaccines and as a method to produce therapeutic neutralizing mAbs that would not be affected by Omicron or future SARS-CoV-2 VOC substitutions.

## Results

### The majority of mosaic-8 elicited mouse mAbs identified as binding two or more RBDs are cross-neutralizing

We made multimerized RBD-nanoparticles using the SpyCatcher-SpyTag system (Brune et al., 2016; Zakeri et al., 2012) to covalently attach RBDs with C-terminal SpyTag003 sequences to a 60-mer nanoparticle (SpyCatcher003-mi3) (Keeble et al., 2019). As previously described, we produced and characterized nanoparticles presenting randomly arranged RBDs from SARS-CoV-2 and seven animal sarbecovirus spikes (mosaic-8 RBD-mi3) as well as nanoparticles presenting only SARS-CoV-2 RBDs (homotypic SARS-CoV-2 RBD-mi3) (Cohen et al., 2021) (Figure S1). Mice were primed and boosted with either mosaic-8 or homotypic SARS-CoV-2 RBD-nanoparticles in AddaVax adjuvant. We used a Berkeley Lights Beacon Optofluidic system to screen a subset of B cells for binding to one or more labeled RBDs (Figure S3). B cells secreting IgGs that bound to at least one RBD were exported from the instrument, and the variable domains of heavy and light chain genes were sequenced and subcloned into expression vectors containing genes encoding human IgG C_H_1-C_H_2-C_H_3 domains, human C_H_1, or human C_L_ domains. From 39 exported cells, we isolated genes for 15 RBD-binding mAbs (Table S1) that were expressed as IgGs and corresponding Fabs. The 15 unique IgG sequences included 13 that were derived from mosaic-8 immunized mice and identified during the screen as binding to two or more labeled RBDs (six mAbs) or to one labeled RBD (seven mAbs), and two were derived from homotypic RBD-nanoparticle immunized mice and identified as binding to two or more labeled RBD (Figure 2A) (Table S1). Two mAbs from mosaic-8 immunized mice were excluded from analyses after showing no detectable binding to purified RBDs (Table S1).

**Figure 2.**
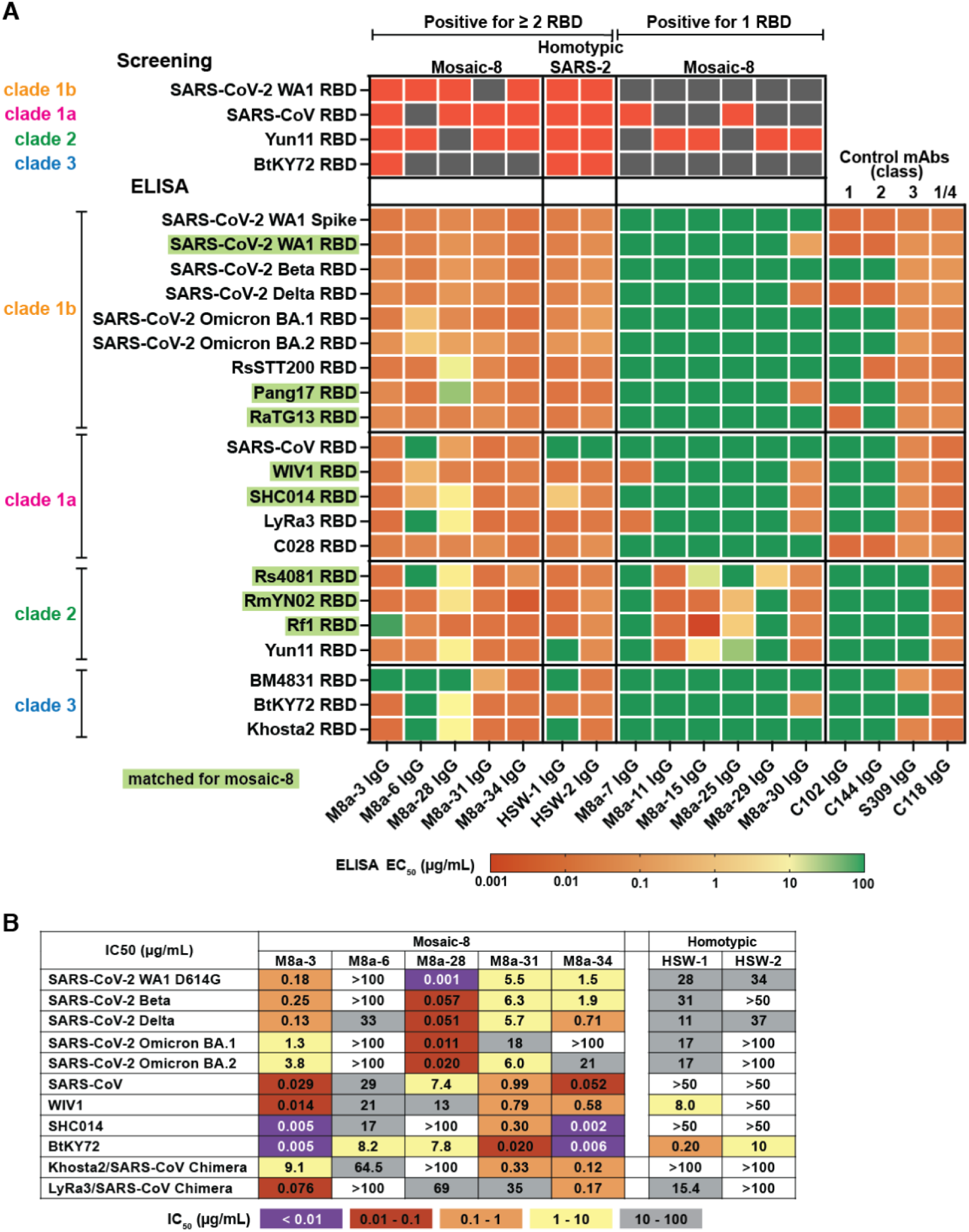
A subset of mAbs elicited in mosaic-8 and homotypic SARS-CoV-2 RBD nanoparticle-immunized mice show cross-reactive binding and neutralization properties. (**A**) Top four rows: RBDs used for screening of single B cells. Red indicates binding; dark gray indicates no binding. Remaining rows: ELISA EC_50_ values for mouse mAb binding to sarbecovirus RBDs from different clades. RBDs that were included on the mosaic-8 RBD-nanoparticles are shaded in green. EC_50_ values were derived from ELISAs conducted with duplicate samples at least twice (for first seven mAbs) or once (for remaining mAbs). M8a-11 and M8a-26 shared the same protein sequences, the same EC_50_ values were presented twice. (**B**) Neutralization potencies (IC_50_ values) of mAbs against SARS-CoV-2 variants and indicated sarbecoviruses. IC_50_ values are reported from neutralization assays that were conducted using duplicate samples at least twice except for a single assay for M8a-28 against Omicron BA.1.

We first evaluated binding of the 13 purified mAb IgGs to RBDs from SARS-CoV-2 variants and other sarbecoviruses using enzyme-linked immunosorbent assays (ELISAs). RBDs were included from sarbecoviruses clades 1a, 1b, 2, and 3 clades (as defined in (Starr et al., 2022)) (Figure 2A). We compared the mAb binding profiles to four human anti-RBD IgGs with known epitopes: C118, a broadly cross-reactive class 1/4 mAb isolated from a convalescent COVID-19 donor (Jette et al., 2021; Robbiani et al., 2020), S309 (Sotrovimab), a cross-reactive class 3 mAb isolated from a SARS-CoV–infected donor (Pinto et al., 2020), and mAbs isolated from COVID-19 donors that bind to more variable RBD epitopes overlapping with the ACE2-binding footprint (Robbiani et al., 2020): C102 (class 1) and C144 (class 2) (Figure 2A). Of the seven mAbs isolated from B cells from vaccinated mice that were identified during screening as secreting IgGs that bound to more than one RBD (Figure 2A), five were isolated from mosaic-8 RBD-nanoparticle-immunized mice (names with M8a prefixes), and two were isolated from homotypic RBD-nanoparticle-immunized mice (names with HSW prefixes). These seven mAbs showed binding to SARS-CoV-2 spike trimer and SARS-CoV-2 RBDs including Beta, Delta, and Omicrons BA.1 and BA.2 in addition to the SARS-CoV-2 WA1 variant included in the mosaic-8 RBD nanoparticles, as well as cross-reactive binding to animal sarbecovirus RBDs (Figure 2A). The half maximal effective concentrations (EC_50_ values) for binding of these mAbs to most of the RBDs ranged from 1 to 10,000 ng/mL (Figure 2A). By comparison, six mAbs isolated from B cells that secreted IgGs that bound only one RBD during screening recognized a smaller subset of RBDs evaluated in our ELISA panel, and none bound to SARS-CoV-2 spike (Figure 2A). Some of these mAbs might prove useful for detecting whether particular RBDs are present on a mosaic RBD-nanoparticle: for example, M8a-7 and M8a-29 could be used to detect WIV1 or Rs4081, respectively, on mosaic-8 RBD-nanoparticles. We proceeded with further analysis of the seven mAbs that were isolated from B cells secreting IgGs that bound to two or more RBDs during screening (Figure 2A).

The five M8a IgGs and two HSW IgGs that showed cross-reactive RBD binding during screening and by ELISA shared amino acid sequence identities of ∼50% to 90% in their V_H_ and V_L_ domains (Figure S4A,B) and had varied lengths for their complementarity-determining regions 3 (CDR3s), which are often critical in antigen recognition (Davies and Metzger, 1983): the mAb CDR3s ranged from 9-16 residues for the heavy chain CDR3 (CDRH3) and all were 9 residues for the light chain CDR3 (CDRL3) (Figure S4C), compared with 11 (IgH) and 9 (Igκ) for average C57Bl/6 mouse antibody CDR3s (Rettig et al., 2018). The CDRH1, CDRH2, and CDRL2 regions were the same lengths across the seven mAbs, whereas the CDRL1 ranged from 6-12 residues (Figure S4). M8a-34 and HSW-1 both had long (16 residues) CDRH3s, and M8a-31 had the shortest (9 residues) CDRH3. By contrast to its shorter than average CDRH3, M8a-31 had the longest CDRL1 (12 residues) compared with M8a-3, M8a-6, M8a-28, and HSW-2, which all included six-residue CDRL1s (Figure S4C). M8a-3 and M8a-6, related by high sequence identities (87.6% for V_H_ and 89.7% for V_L_) (Figure S4B) and the shared V gene segments (IgH V1-69 and Igκ V6-25) (Figure S4A, Table S1), both contained 14-residue CDRH3s and six-residue CDRL1s (Figure S4C), yet M8a-3 showed a broader RBD binding profile by ELISA, such that it bound all RBDs evaluated except for the clade 2 Rf1 and clade 3 BM4831 RBDs, whereas M8a-6 did not bind detectably to any of the three clade 3 RBDs or to three of the clade 1a and clade 2 RBDs (Figure 2A). M8a-28 showed weak binding to some non-SARS-2 RBDs of clade 1b (RsSTT200 and Pang17), clade 1a (SHC014 and LyRa3) and clade 2 (Rs4081, RmYN02 and Yun11), and weak or no binding to RBDs of clade 3 (weak for BtKY72 and Khosta-2, and no binding to BM4831 RBD of clade 3 (Figure 2A). In contrast, HSW-2 showed binding to RBDs from all clades except SARS-CoV from clade 1a (Figure 2A). M8a-31 and M8a-34 recognized all RBDs in the ELISA panel (Figure 2A). Although M8a-34 and HSW-1 shared a sequence identity of 75.3% for V_H_ and 88.3% for V_L_ with the same light chain IgκV3-5 V gene segment (Figure S4A, Table S1), and both had 16-residue CDRH3s and 10-residue CDRL1s (Figure S4B,C), HSW-1 was not as broadly cross-reactive by ELISA, showing no detectable binding to RBDs of SARS-CoV (clade 1a), Yun 11 (clade 2), or BM4831 and Khosta2 (clade 3) (Figure 2A).

We next measured neutralization potencies using a pseudovirus neutralization assay (Crawford et al., 2020) against sarbecoviruses known to use human ACE2 for target cell entry, including SARS-CoV-2 WA1 D614G, SARS-CoV-2 Beta, SARS-CoV-2 Delta, SARS-CoV-2 Omicron BA.1 and BA.2, SARS-CoV, WIV1, SHC014, a modified BtKY72 (Starr et al., 2022) Khosta2/SARS-CoV Chimera, and LyRa3/SARS-CoV Chimera (Figure 2B). Among all, M8a-3 was the most consistently cross-reactively potent, with low half-maximal inhibitory concentrations (IC_50_ values) against all pseudoviruses evaluated (Figure 2B). Despite sharing high sequence identity, the same V gene segments, and similar CDR characteristics with M8a-3 (Figure S4), M8a-6 showed no neutralizing activity except for weak activity against SHC014 and BtKY72. A less related mAb, M8a-28, was a potent neutralizer, but only against SARS-CoV-2 variants. The M8a-31 and M8a-34 mAbs were less potent against SARS-CoV-2 variants, but were more broadly cross-reactive, correlating with their ELISA profiles (Figure 2B). By contrast to the five M8a mAbs, the HSW-1 and HSW-2 mAbs isolated from homotypic SARS-CoV-2 RBD nanoparticle-immunized mice identified as binding two or more RBDs during screening showed overall weaker neutralizing potencies, with eight of 18 assays showing no neutralizing activity and most of the remaining assays showing IC_50_ values above 10 µg/mL (Figure 2B).

To identify RBD epitopes for the mAbs, we used a binding assay to assess potential competition with proteins that bind to known RBD epitopes. For this competition assay, we used the four human anti-RBD mAbs with known epitopes that were used as controls for ELISAs (Figure 2A): C118 (class 1/4), S309 (class 3), C102 (class 1), and C144 (class 2), along with other potential competitor mAbs or control mAbs: C022 (class 1/4) (Jette et al., 2021; Robbiani et al., 2020), CR3022 (class 4) (Huo et al., 2020), COVA1-16 (Liu et al., 2020a), C135 (class 3), C110 (class 3), C105 (class 1) (Robbiani et al., 2020), and a soluble human ACE2-Fc construct (Jette et al., 2021). The ELISA revealed the expected competition for the characterized human mAbs (Figure S5), validating its use for mapping RBD epitopes for the seven mouse mAbs elicited by RBD-nanoparticle immunization. Three of the five m8a mAbs (M8a-3, M8a-31, and M8a-34) mapped to class 1/4 and/or class 4 epitopes, M8a-28 mapped to the class 3 RBD region, and Ma-6 did not compete with any of the labeled anti-RBD IgGs. The identification of a class 3 RBD epitope for M8a-28 rationalized its potent neutralization of SARS-CoV-2 WA1 and VOCs and limited cross-reactive neutralization of animal sarbecoviruses (Figure 2B), by contrast with the class 1/4 RBD epitope identification for the remaining less potently neutralizing M8a mAbs (M8a-3, M8a-31, and M8a-34), since this class of anti-RBD mAb tends to show less potent neutralization, but broader sarbecovirus cross-reactivity, than other classes due to the more occluded nature of the class 1/4 epitope (Cohen et al., 2022; Jette et al., 2021; Tortorici et al., 2021). Of the two HSW mAbs, HSW-1 did not exhibit any detectable competition, and HSW-2 showed competition with CR3022, a class 4 anti-RBD mAb. Overall, these results demonstrated that the majority of the mAbs identified during Beacon screening mapped to the more conserved class 1/4, 4, and 3 RBD epitopes.

### Cryo-EM structures of Fab-spike trimer complexes reveal cross-reactive recognition and rationalize neutralization results

To deduce recognition and neutralization mechanisms, we performed structural analyses of the seven mAbs identified as binding two or more RBDs during the Beacon isolation (Figure 2A). Single-particle cryo-EM was used to solve structures of complexes of a SARS-CoV-2 6P spike trimer (Hsieh et al., 2020) and a mAb Fab, five of which were derived from M8a IgGs isolated in mosaic-8 nanoparticle–immunized mice (Figure 3, Figure 4A) and two from HSW IgGs elicited in homotypic SARS-CoV-2 RBD nanoparticle–immunized mice (Figure 5) (Figure S6-13, Table S2).

**Figure 3.**
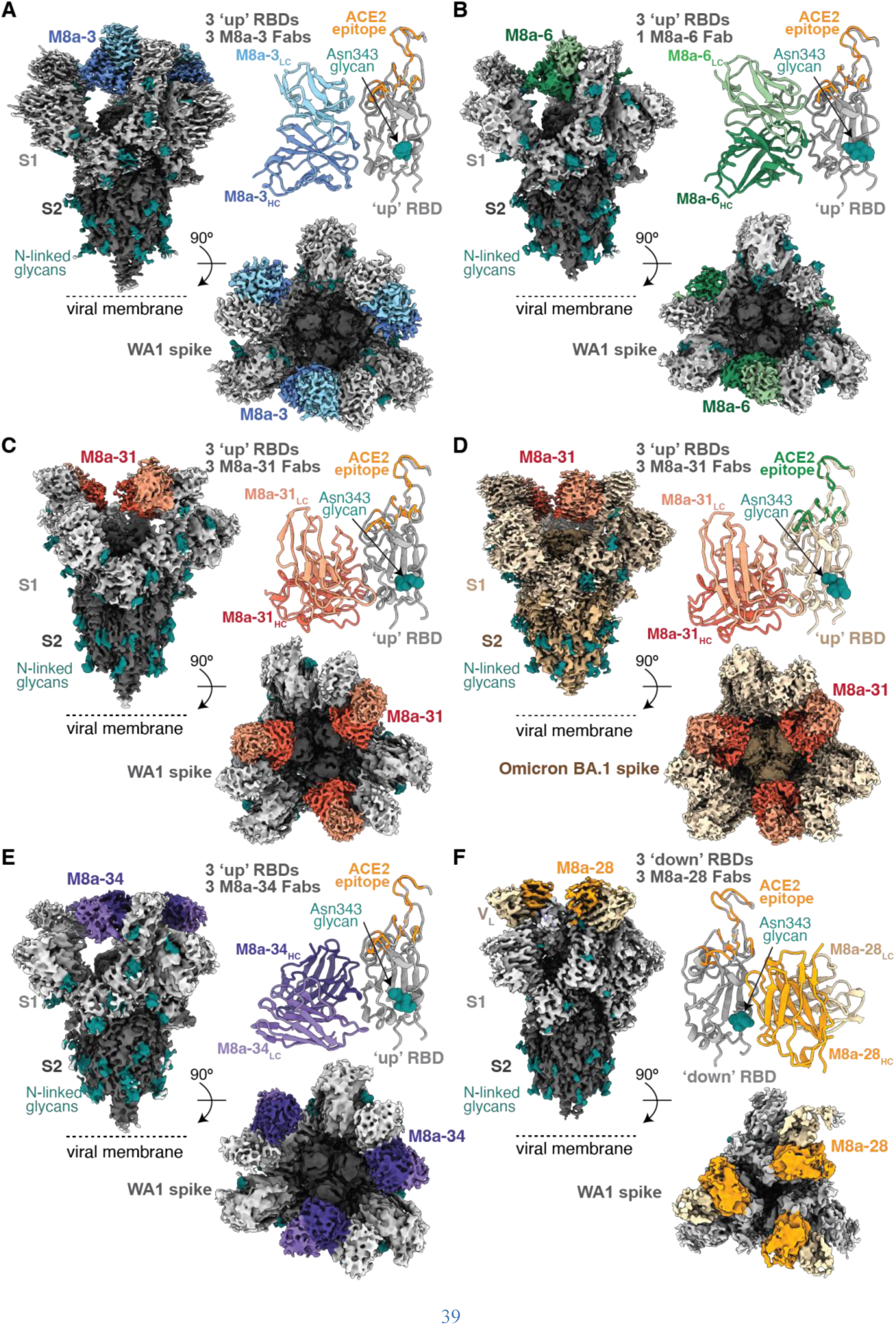
mAbs isolated from mice immunized with mosaic-8 nanoparticles target epitopes outside of the ACE2 binding footprint. EM densities of single-particle cryo-EM structures of Fab-spike trimer complexes are shown from the side (upper left), top (lower right), and as cartoon diagrams of the Fab V_H_-V_L_ interaction with the RBD (upper right; RBD residues involved in ACE2 binding are in colored in orange for complexes with WA1 spike and in green for complex with Omicron BA.1 spike). Only V_H_-V_L_ domains are shown for each Fab. (**A**) WA1 spike complexed with M8a-3. (**B**) WA1 spike complexed with M8a-6. (**C**) WA1 spike complexed with M8a-31. (**D**) Omicron BA.1 spike complexed with M8a-31. (**E**) WA1 spike complexed with M8a-34. (**F**) WA1 spike complexed with M8a-28.

**Figure 4.**
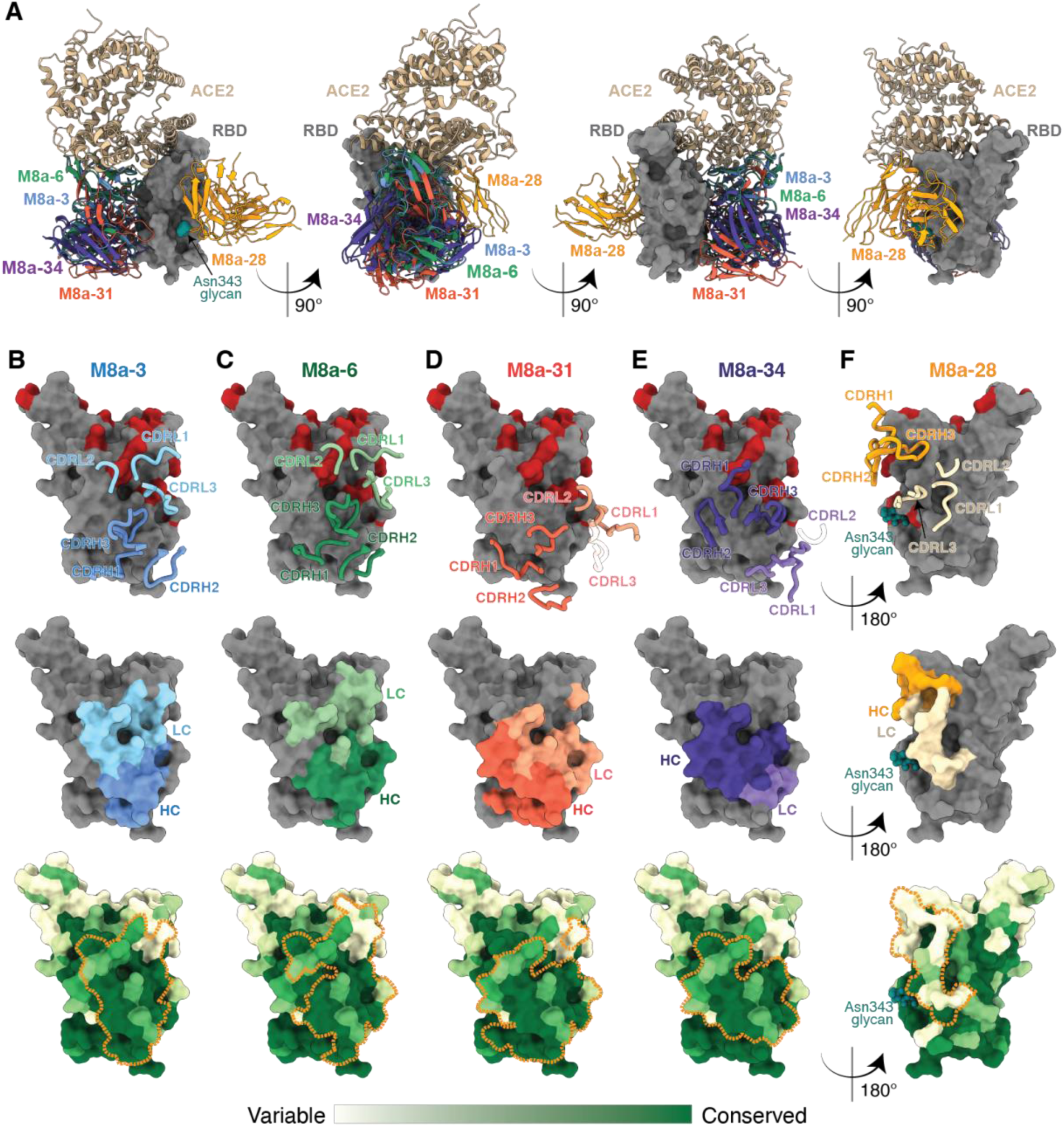
Epitopes of mAbs elicited by mosaic-8 immunization demonstrate targeting of non-class 1/class 2 RBD epitopes. (**A**) Four views of RBD surface (dark gray) with overlay of mAb V_H_-V_L_ domains (different colored cartoon representations) from Fab-spike structures. ACE2 (tan cartoon representation) complexed with RBD (PDB 6M0J) is shown for comparison. **(B-F**) mAb epitopes on RBD surface shown with overlaid heavy and light chain CDRs (top, CDRs that do not interact with the RBD are shown in transparent cartoons; all CDRs were defined based on the IMGT definition), as colored areas for heavy and light chains (middle) and outlined with orange dotted lines on a sequence conservation surface plot (bottom; calculated using the 16 sarbecovirus RBD sequences shown in Figure S1). The N-glycan at RBD position Asn343 was shown as spheres. (**B**) M8a-3. (**C**) M8a-6. (**D**) M8a-31 from complex with WA1 spike trimer. (**E**) M8a-34. (**F**) M8a-28. Omicron BA.1 and BA.2 substitutions were colored red in the top panels of (**B**-**F**).

**Figure 5.**
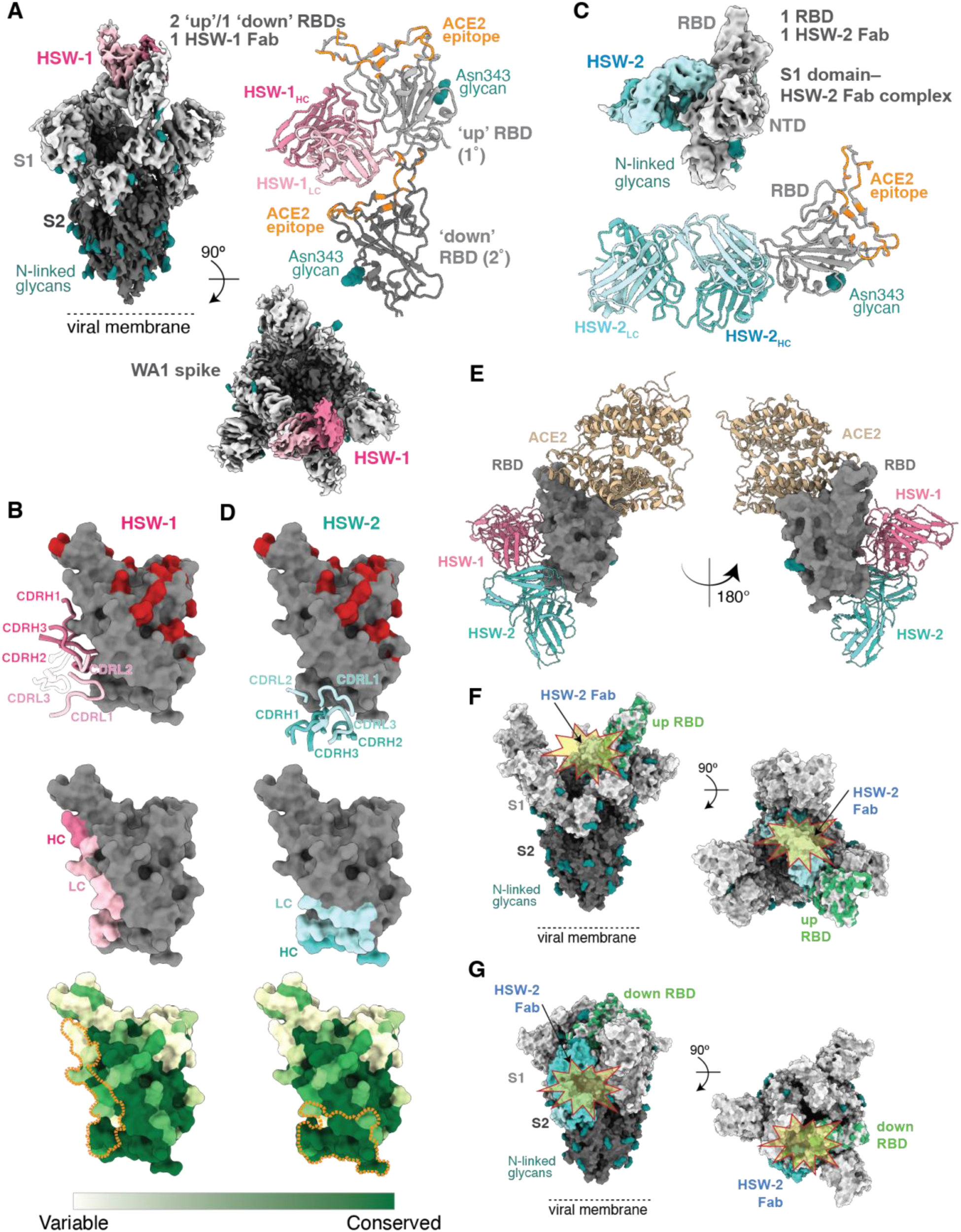
Epitopes of mAbs isolated from mice immunized with homotypic SARS-CoV-2 nanoparticles that target conserved RBD epitopes. (**A**) EM density of single-particle cryo-EM structure of HSW-1 Fab–spike trimer complex shown from the side (upper left), the top (lower right), and as a cartoon diagram of the HSW-1 V_H_-V_L_ interaction with two adjacent RBDs (1° and 2°) (upper right). HSW-1 interacts mainly with an ‘up’ RBD (1° RBD, light grey) but also includes V_L_ interactions with a ‘down’ RBD (2° RBD, dark grey). (**B**) HSW-1 epitope on RBD surface shown with overlaid heavy and light chain CDRs (top, CDRs that do not interact with the RBD are shown in transparent cartoons; all CDRs were defined based on the IMGT definition), as colored areas for heavy and light chains (middle) and outlined with orange dotted lines on a sequence conservation surface plot (bottom; calculated using the 16 sarbecovirus RBD sequences in Figure S1). Omicron BA.1 and BA.2 substitutions were colored red in the top panel. (**C**) EM density of single-particle cryo-EM structure of HSW-2–Fab S1 domain complex (top) and cartoon diagram of the HSW-2 V_H_-V_L_ interaction with the RBD (bottom). (**D**) HSW-1 epitope on RBD surface shown with overlaid heavy and light chain CDRs (top, all CDRs were defined based on the IMGT definition), as colored areas for heavy and light chains (middle) and outlined with orange dotted lines on a sequence conservation surface plot (bottom; calculated using the 16 sarbecovirus RBD sequences shown in Figure S1). Omicron BA.1 and BA.2 substitutions were colored red in the top panel. (**E**) Two views of RBD surface (dark gray) with overlays of mAb V_H_-V_L_ domains (different colored cartoon representations) from HSW Fab-spike structures and ACE2 (tan cartoon representation from PDB 6M0J). (**F,G**) Superpositions of HSW-2–RBD structure onto the RBD from a spike trimer structure showing that HSW-2 Fab is sterically hindered from binding to either an ‘up’ or ‘down’ RBD on an intact spike trimer due to clashes (indicated as starbursts) with the spike S2 domain. (**F**) HSW-2 Fab-RBD interaction modeled onto an ‘up’ RBD from the M8a-31– spike complex structure. (**G**) HSW-2 Fab-RBD interaction modeled onto a ‘down’ RBD from the M8a-28–spike complex structure.

Each of the five M8a mAb Fab structures were solved as complexes with the SARS-CoV-2 WA1 spike trimer, and the M8a-31 Fab was also solved complexed with the Omicron BA.1 spike (Figure 3A-F; Figure S6-S11; Table S2). For five of the six M8a-spike structures (WA1 spike plus Fabs from M8a-3, M8a-31, M8a-34, M8a-28 and Omicron BA.1 spike plus M8a-31 Fab), we observed one Fab bound to each of the three RBDs, which were in ‘up’ positions in all cases except for the M8a-28–spike structure in which all three RBDs were in the ‘down’ conformation (Figure 3C). In the remaining structure (M8a-6–spike), we found only one well-resolved Fab per trimer.

A 3.1 Å resolution M8a-3 Fab–spike complex structure revealed Fab V_H_-V_L_ interactions with ‘up’ RBDs using all six CDRs along with residues within the light chain framework region 2 and 3 (FWRL2 and FWRL3) (Figure 3A, 4B, Figure S6, S14A). Consistent with the competition ELISA results (Figure S5), comparison of the M8a-3 Fab-RBD interaction with previously-characterized representative anti-RBD antibodies in different structural classes (Barnes et al., 2020a; Jette et al., 2021) showed overlap with the class 1 and class 4 RBD epitopes (Figure S14A) and a binding footprint adjacent to that of ACE2 (Figure 3A, 4A), similar to the human mAb C118, a class 1/4 anti-RBD antibody that blocks ACE2 binding without substantially overlapping with the ACE2 binding footprint (Jette et al., 2021) and that competes for binding to RBD with M8a-3 (Figure S5). The M8a-3–spike structure showing recognition of a largely conserved region of the RBD (Figure 4B) was consistent with ELISA and neutralization results demonstrating that M8a-3 neutralized and/or bound to most of the sarbecoviruses and the SARS-CoV-2 variants tested (Figure 2).

A 3.2 Å spike trimer structure complexes with the related, but mostly non-neutralizing M8a-6 mAb, showed three ‘up” RBDs, but only one well-resolved Fab (Figure 3B; Figure S7, S14B), which we modeled using information from a partially-refined 3.0 Å M8a-6 Fab–RBD crystal structure. The M8a-6 Fab shared a similar RBD binding epitope and approach angle as M8a-3 (Figure 3A, 4A, Figure S14B), interacting with the RBD using all six CDRs plus framework regions FWRH2, FWRL2, and FWRL3 (Figure 4C). Furthermore, M8a-6 also recognized a similar epitope as the C118 (Jette et al., 2021) and M8a-3 mAbs, involving mostly conserved RBD residues (Figure 4C, Figure S14B). Despite sharing high sequence identity and similar binding epitopes on SARS-CoV-2 RBD with M8a-3, M8a-6 was non-neutralizing against SARS-CoV-2 and only weakly neutralizing against SHC014, whereas M8a-3 neutralized SARS-CoV-2 D614G with a 0.18 µg/mL IC_50_ (Figure 2B). These different neutralization profiles likely result from a weaker interaction of M8a-6 as compared with M8a-3 with CoV spikes, as demonstrated by incomplete binding of Fabs in the M8a-6–spike complex cryo-EM structure and the lack of competition of M8a-6 IgG with any of the biotinylated IgGs, including C118, with known epitopes (Figure S5).

In common with M8a-3, M8a-31 exhibited broadly cross-reactive binding and neutralization across SARS-CoV-2 variants and other sarbecoviruses (Figure 2) and competed with class 1/4 and class 4 anti-RBD antibodies (Jette et al., 2021) (Figure S5). Single-particle cryo-EM structures were determined for M8a-31 Fab bound to SARS-CoV-2 WA1 (Figure 3C, Figure S8) and to Omicron BA.1 (Figure 3D, Figure S9) spike trimers at resolutions of 2.9 Å and 3.5 Å, respectively. In both structures, three M8a-31 Fabs interacted with ‘up’ RBDs (Figure 3C,D, Figure S14C,D). Despite the 15 substitutions in the Omicron BA.1 RBD compared with the WA1 RBD, the epitope and binding pose of the M8a-31 Fab in both structures were similar (Figure S14C,D) with a root mean square deviation (RMSD) of 1.0 Å (calculated using all 1,267 resolved Cα atoms in each Fab-spike protomer structure). The binding of M8a-31 Fab to both the SARS-CoV-2 WA1 and Omicron BA.1 RBDs was mainly stabilized through interactions with all CDRs except for CDRL3 along with FWRH1, FWRH2, FWRL2, and FWRL3 (Figure 4D). The M8a-31 epitope overlapped with class 4 anti-RBD antibodies, but was shifted towards the ACE2 binding site compared with the CR3022 class 4 mAb (Figure S14C,D), consistent with its competition with the C118 class 1/4 mAb (Figure S5). The mainly conserved nature of the M8a-31 epitope (Figure 4D) is consistent with its broadly cross-reactive binding and neutralization properties (Figure 2).

In common with M8a-3 and M8a-31, M8a-34 bound and neutralized most sarbecoviruses across different clades and SARS-CoV-2 variants (Figure 2) and exhibited a similar competition profile (Figure S5). To map its epitope, we determined a single-particle cryo-EM structure of M8a-34 Fab bound to the SARS-CoV-2 WA1 spike trimer at 3.5 Å resolution (Figure 3E, Figure S10), revealing interactions of three Fabs with three ‘up’ RBDs (Figure 3E, Figure S14E) that were modeled using an M8a-34 Fab–RBD crystal structure (Table S3). M8a-34 Fab interacted with the RBD through all three CDRHs as well as CDRL1 and CDRL3 (Figure 4E). The M8a-34 Fab epitope was similar to epitopes of other class 1/4 mAbs including M8a-3, M8a-6 and M8a-31, which overlapped with the binding epitopes of CR3022 (class 4) and C118 (class 1/4) (Figure 4A, Figure S14E), again consistent with its broad binding and neutralizing properties (Figure 2) and competition ELISA results (Figure S5).

The M8a-28 mAb showed the lowest degree of cross-reactive RBD binding (Figure 2A). M8a-28 mapped to the class 3 epitope instead of the more conserved class 1/4 and class 4 epitopes (Figure S5), and except for M8a-6 (a weakly/non-neutralizing mAb), it showed the lowest levels of cross-reactive sarbecovirus neutralization of the five mAbs isolated from mosaic-8 immunized mice (Figure 2B). Single-particle cryo-EM structures of the M8a-28 Fab in complex with the SARS-CoV-2 spike were determined in two conformational states: a 2.8 Å structure with each of three Fabs binding to a ‘down’ RBD (Figure 3F) and a 3.1 Å structure with two Fabs bound to adjacent ‘down’ RBDs and a third Fab at lower occupancy bound to a flexible ‘up’ RBD (Figure S11). The Fab-RBD interaction was mediated by all six CDRs, plus FWRH3 and FWRL1 (Figure 4F). Notably, the M8a-28 Fab approached the RBD from the opposite direction compared with Fabs from the other M8a mAbs (Figure 4A, Figure S14F), interacting with more variable RBD regions (Figure 4F) that overlapped with the epitope of the class 3 anti-RBD mAb, S309 (Pinto et al., 2020) (Figure S14F). Although M8a-28 potently neutralized SARS-CoV-2 WA1 D614G, Beta, Delta, Omicron BA.1 and BA.2, it was only weakly neutralizing or non-neutralizing against other sarbecoviruses (Figure 2B), consistent with its epitope spanning more variable RBD residues than epitopes of class 4 and class 1/4 anti-RBD mAbs (Jette et al., 2021).

The HSW-1 and HSW-2 mAbs, isolated from homotypic SARS-CoV-2 RBD nanoparticle-immunized mice, were each isolated from B cells secreting IgGs that bound to all four RBDs used during screening. Accordingly, they exhibited broad cross-reactive binding to sarbecovirus RBDs by ELISA, with HSW-2 binding to all tested RBDs except for SARS-CoV RBD and HSW-1 also not interacting detectably with SARS-CoV-1 RBD and three other RBDs from clade 2 and 3 sarbecoviruses (Figure 2A). Despite broad recognition of sarbecovirus RBDs, the HSW mAbs especially HSW-2, exhibited overall weaker neutralization potencies across the sarbecoviruses tested, with all IC_50_ values being above 10 µg/mL (Figure 2B). To compare recognition properties with the M8a Fabs, we determined a single-particle cryo-EM structure of HSW-1 bound to SARS-CoV-2 WA1 spike trimer at 3.1 Å resolution, revealing a single well-ordered Fab bound to a trimer with two ‘up’ RBDs and one ‘down’ RBD (Figure 5A, Figure S12, Figure S15A). In common with the M8a-6 mAb, for which the Fab-spike structure also revealed only a single bound Fab per trimer (Figure 3B), HSW-1 showed no detectable competition with anti-RBD mAbs with known epitopes (Figure S5). The bound HSW-1 Fab interacted with two RBDs: one ‘up’ RBD (1° RBD) and the adjacent ‘down’ RBD (2° RBD) (Figure 5A, Figure S15A). Interactions between the HSW-1 Fab and 1° RBD were mediated by both the heavy chain through FWRH1, CDRH1 and CDRH3, and the light chain through CDRL1, CDRL2, CDRL3 and FWRL2 (Figure 5A,B). The interactions between HSW-1 and the 2° RBD were mediated by the HSW-1 light chain (Figure 5A). Structural comparisons showed the epitope of HSW-1 overlapped somewhat with the binding epitopes of C118 (class 1/4) and CR3022 (class 4) and included mostly conserved residues (Figure S15A).

We next used single-particle cryo-EM to investigate the interactions of HSW-2 with spike trimer. We observed two main populations of particles, one population of unliganded intact spike trimer and a second HSW-2 Fab complexed with spike S1 domain protomers (Figure S13). From the second population, we obtained an EM reconstruction at 4.1 Å resolution of the HSW-2 Fab bound to the S1 domain of the SARS-CoV-2 WA1 spike (Figure 5C, Figure S15B) using a crystal structure of an HSW-2 Fab–RBD complex (Table S3) to derive detailed interactions. HSW-2 used its six CDRs plus FWRH2, FWRL1, FWRL2 and FWRL3 to recognize the bottom of the RBD (Figure 5D,E), consistent with its competition with the class 4 anti-RBD antibody CR3022 (Figure S5). Although their binding poses differed, the HSW-2 epitope overlapped with the largely-conserved epitope of the class 4 mAb, CR3022 (Figure S15B), a mAb isolated from a SARS-CoV patient that also induces dissociation of the SARS-CoV-2 spike trimer (Huo et al., 2020). Although S1 shedding resulting from mAb binding has been suggested as a possible neutralization mechanism for CR3022 and other class 4 anti-RBD mAbs (Huo et al., 2020; Piccoli et al., 2020; Wec et al., 2020), HSW-2 was largely non-neutralizing (Figure 2B). To determine whether HSW-2 was raised against an RBD epitope that is inaccessible in an intact spike trimer, we aligned the RBD portion of the HSW-2 Fab-RBD structure to RBDs from a spike trimer structure with all ‘up’ RBDs and to a spike structure with all ‘down’ RBDs, finding steric clashes in both cases (Figure 5F,G). The inability of the HSW-2 Fab to access either ‘up’ or ‘down’ RBDs in the context of an intact spike trimer is consistent with the observation that although HSW-2 bound to almost all RBDs evaluated by ELISA (Figure 2A), it showed weak or no neutralization activity against the tested strains (Figure 2B).

### Class 1/4 anti-RBD mAbs induce spike trimer opening and exhibit different potentials for intra-spike crosslinking and susceptibility to mutations

To address potential effects of mAb binding on spike trimer conformation, we compared the Fab-bound spike trimer structures reported here to other trimer structures. We previously used the distances between the Cα atoms of residue 428 in each of three ‘up’ RBDs within an intact trimer to assess spike openness, with inter-protomer distances of 39 Å or less indicating a typical prefusion spike trimer conformation (Figure 6A) (either unliganded, bound to ACE2, or bound to a class 1, 2, or 3 anti-RBD mAb), and binding of class 4 and class 1/4 anti-RBD mAbs resulting in increases in the inter-RBD distances (Jette et al., 2021). For example, inter-RBD distances for spike trimers bound to C118 or to S2X259, human donor-derived mAbs recognizing class 1/4 epitopes (Jette et al., 2021; Robbiani et al., 2020; Tortorici et al., 2021), were 53 Å (Jette et al., 2021) (C118) or 43 Å (S2X259), demonstrating opening of the trimer to accommodate a Fab interacting with the occluded class 1/4 region of the RBD (Figure 6B). In the present study, we found inter-protomer distances ranging from 48-69 Å for trimers bound to Fabs from M8a-3 (Figure 6C), M8a-6 (Figure 6D), M8a-31 (Figure 6E,F), M8a-34 (Figure 6G) and HSW-1 (Figure f16H), consistent with increased openness of trimers bound to class 1/4 and class 4 anti-RBD antibodies. By contrast, the comparable inter-protomer distance was 31 Å in the M8a-28–spike structure with all RBDs in the ‘down’ conformation (Figure 6I), consistent with M8a-28 recognition of the non-occluded class 3 RBD epitope.

**Figure 6.**
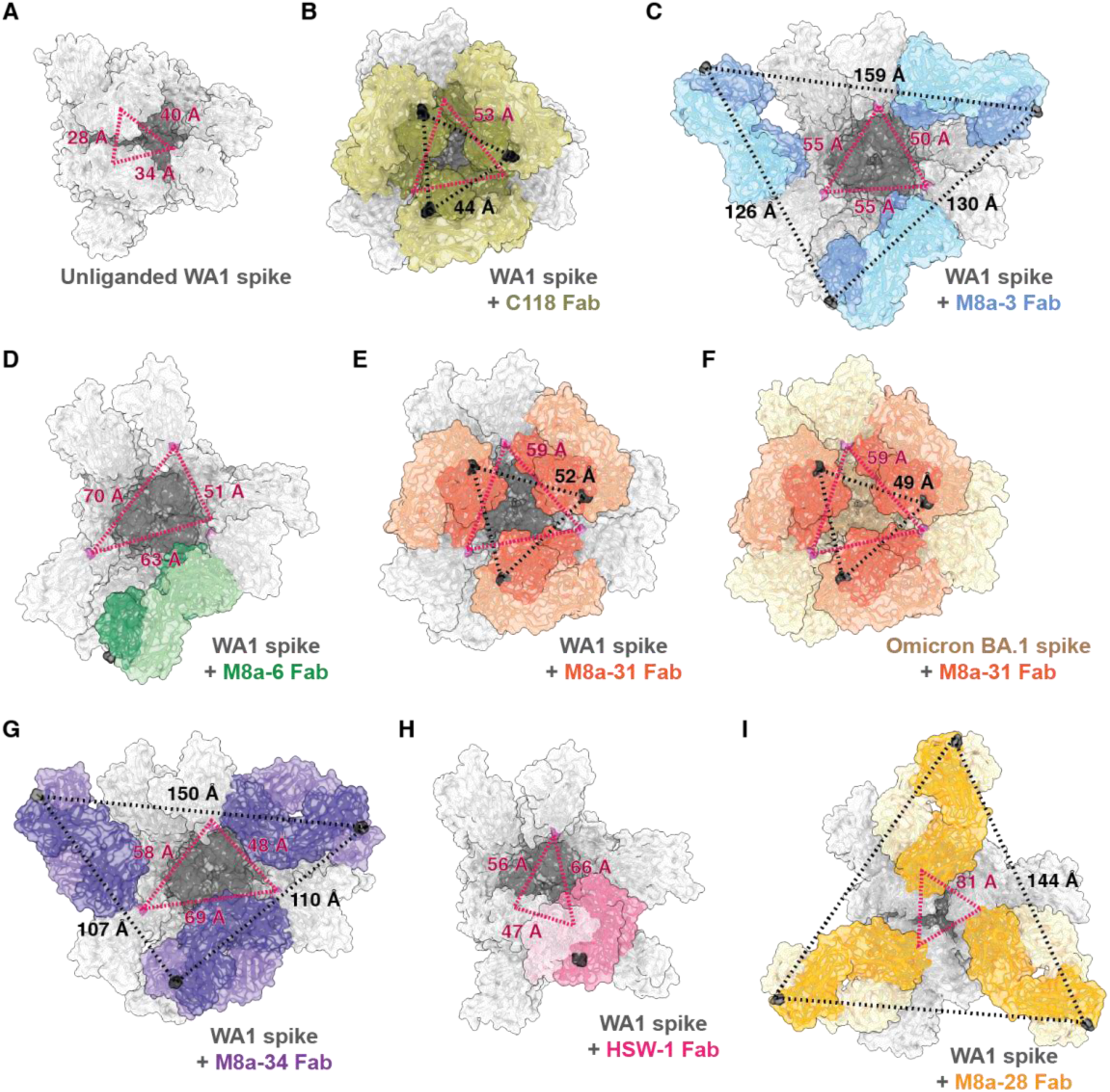
Spike-mAb complex structures show increased trimer openness and the potential for intra-spike IgG crosslinking. Red dotted lines: Trimer openness was assessed by measuring distances between the Cα atoms of RBD residue 428 (pink) in each RBD of a spike trimer from the indicated Fab-spike complex structures (top-down views with mAb Fabs shown in colors on a gray spike trimer (WA1) or an orange spike trimer (Omicron BA.1). Distances of 39 Å or less indicate a typical closed, prefusion spike trimer conformation (Barnes et al., 2020a) (panel A). Binding of class 1/4 anti-RBD antibodies such as C118 and S2X259 result in larger inter-RBD distances: 53 Å for C118 (panel B) and 43 Å for S2X259 (PDB 7RA8), indicating a more open trimer conformation. Black dotted lines: The potential for intra-spike crosslinking by the two Fabs of a single bound IgG was assessed by measuring distances between the Cα atoms of C-terminal C_H_1 residues (black) on adjacent bound Fabs on the RBDs of a spike trimer. Distances less than 65 Å are considered compatible with the potential for intra-spike crosslinking (Barnes et al., 2020a). (**A**) Unliganded spike (PDB 6VYB): closed prefusion conformation. (**B**) C118 Fab–WA1 spike (PDB 7RKV): open trimer conformation with potential for intra-spike crosslinking by C118 IgG. (**C**) M8a-3 Fab–WA1 spike: open trimer confirmation with no potential for intra-spike crosslinking. (**D**) M8a-6–WA1 spike: open trimer conformation. Black dotted lines between the Cα atoms of C-terminal C_H_1 residues are not shown because the reconstruction included only one Fab. (**E**) M8a-31–WA1 spike: open trimer conformation with potential for intra-spike crosslinking by M8a-31 IgG. (**F**) M8a-31–Omicron BA.1 spike: open trimer conformation with potential for intra-spike crosslinking by M8a-31 IgG. (**G**) M8a-34–WA1 spike: open trimer conformation with no potential for intra-spike crosslinking by M8a-34 IgG. (**H**) HSW-1–WA1 spike: open trimer conformation. Black dotted lines between the Cα atoms of C-terminal C_H_1 residues are not shown because the reconstruction included only one Fab. (**I**) M8a-28–WA1 spike: closed trimer conformation with no potential for intra-spike crosslinking.

To understand how substitutions in recent VOCs might affect binding of the mAbs for which we had Fab-spike structures, we mapped their binding epitopes compared to the locations of Omicron BA.1 and BA.2 substitutions on the RBD (Figure 4B-F, 5B,D). Most of the Omicron BA.1 an BA.2 substitutions were in the ACE2 binding region (Figure 4A, Figure S2B), which includes mainly variable residues (Figure 1A), with fewer substitutions in conserved regions (Figure 1A, 4B-F, 5B,D, Figure S2B). The Omicron substitutions were mainly at the peripheries of the RBD epitopes of the m8a mAbs isolated from mosaic-8 nanoparticle-immunized mice (Figure 4B-F), and there were no Omicron BA.1 or BA.2 substitutions within the binding epitopes of the two HSW mAbs isolated from homotypic SARS-CoV-2 nanoparticle-immunized mice (Figure 5B,D). Despite the Omicron BA.1 or BA.2 substitutions not greatly affecting RBD binding by the seven mAbs (Figure 2A), some of the class 1/4 M8a mAbs showed somewhat reduced neutralization potencies (Figure 2B). Evaluation of potent class 1, 2 and 3 anti-RBD human mAbs also revealed that, except for LY-CoV1404 (Bebtelovimab), most showed diminished binding and neutralizing activities against Omicron BA.1 (Zhou et al., 2022). Epitope analysis showed that LY-CoV1404 targets a class 3 RBD epitope adjacent to Omicron BA.1 and BA.2 mutations that is only somewhat conserved with respect to variability within sarbecoviruses, but unlike other EUA-approved class 3 anti-RBD therapeutic mAbs, it was minimally impacted by those substitutions (Figure 7).

**Figure 7.**
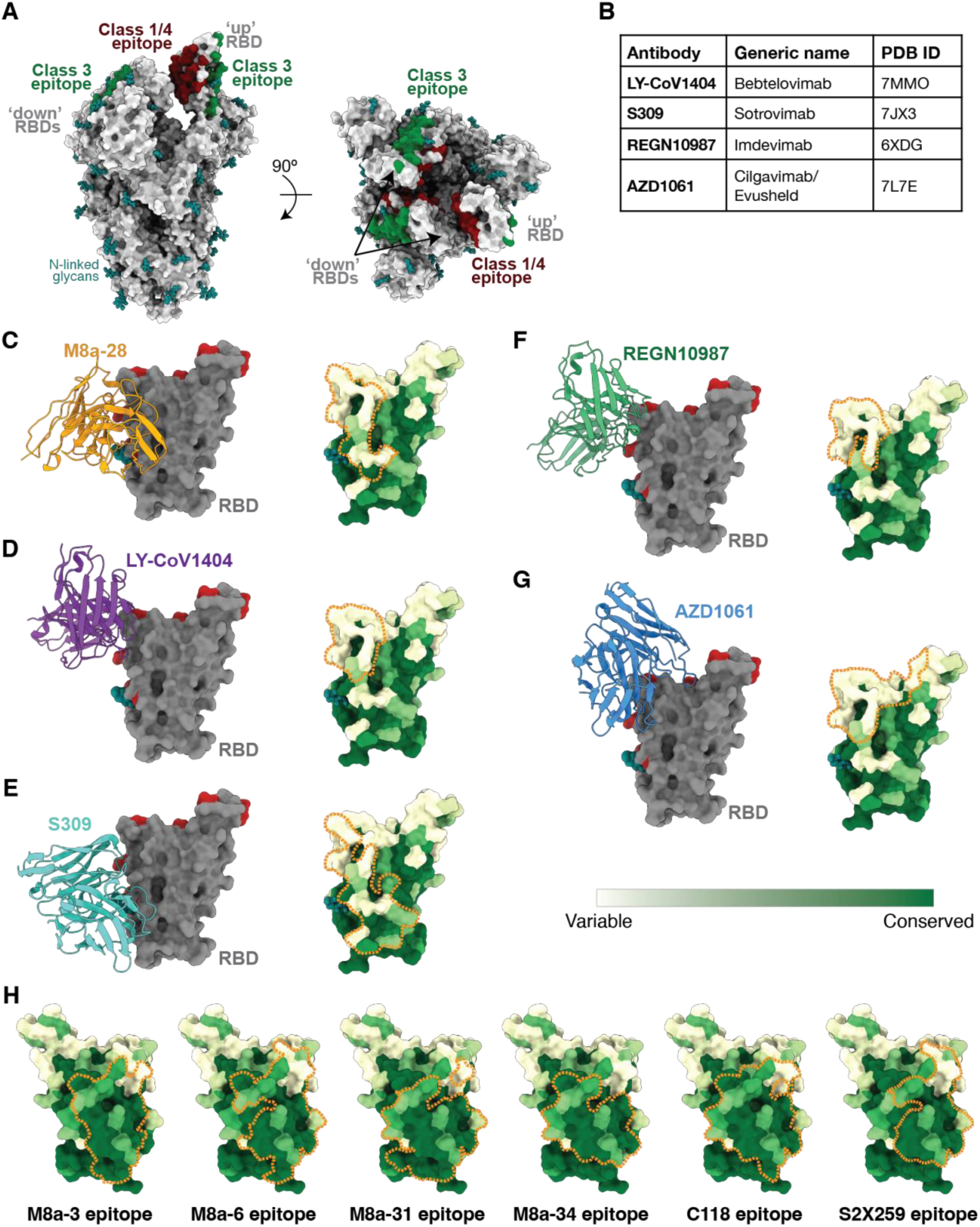
Comparison of M8a epitopes with human mAbs targeting class 3 or class 1/4 RBD epitopes. (**A**) Locations of class 3 and class 1/4 RBD epitopes mapped on an unliganded spike trimer structure with two ‘down’ and one ‘up’ RBDs (PDB 6VYB) showing that the class 3 epitope is exposed, whereas the class 1/4 epitope is partially occluded in the context of the spike trimer. The binding epitopes of representative class 3 (S309/Sotrovimab, PDB 7JX3) and class 1/4 (C118, PDB 7RKV) anti-RBD antibodies were identified by PDBePISA (Krissinel and Henrick, 2007). (**B**) Class 3 anti-RBD mAbs that currently or previously received Emergency Use Authorization (EUA) approval for human administration by the US Food and Drug Administration (modified from (Zhou et al., 2022)). Of these mAbs, only LY-CoV1404/Bebtelovimab retains full neutralization potency against Omicron BA.1 (Zhou et al., 2022), and the NIH COVID-19 treatment guidelines recommend against use of Bamlanivimab plus Etesevimab, Casirivimab plus Imdevimab, or Sotrovimab for the treatment of COVID-19 (US Food and Drug Administration fact sheets listed below). (**C-G**) Left: V_H_-V_L_ domains of M8a-28 and currently or previously EUA-approved class 3 anti-RBD mAbs (cartoon representations) shown interacting with an RBD (gray surface representation with Omicron BA.1 and BA.2 substitutions in red and the RBD Asn343 N-glycan shown as teal spheres). Right: mAb epitopes outlined with orange dotted lines on a sequence conservation surface plot (calculated using the 16 sarbecovirus RBD sequences shown in Figure S1). (**C**) M8a-28. (**D**) LY-CoV1404/Bebtelovimab (PDB 7MMO). (**E**) S309/Sotrovimab (PDB 7JX3). (**F**) REGN10987/Imdevimab (PDB 6XDG). (**G**) AZD1061/Cilgavimab (PDB 7L7E). (**H**) Comparison of the class 1/4 epitopes of M8a mouse mAbs isolated in these studies with the epitopes of C118 (PDB 7RKV), a broadly cross-reactive human mAb (Jette et al., 2021; Robbiani et al., 2020), and S2X259 (PDB 7RAL), a broadly reactive and potent human class 1/4 mAb (Tortorici et al., 2021). Food and Drug Administration. Fact sheet for healthcare providers: emergency use authorization (EUA) of sotrovimab. 2022. Available at: https://www.fda.gov/media/149534/download. Food and Drug Administration. Fact sheet for healthcare providers: emergency use authorization for Evusheld (tixagevimab co-packaged with cilgavimab). 2022. Available at: https://www.fda.gov/media/154701/download. Food and Drug Administration. Fact sheet for health care providers: emergency use authorization (EUA) of bamlanivimab and etesevimab. 2022. Available at: https://www.fda.gov/media/145802/download. Food and Drug Administration. Fact sheet for health care providers: emergency use authorization (EUA) of REGEN-COV (casirivimab and imdevimab). 2021. Available at: https://www.fda.gov/media/145611/download.

Although we found that RBD binding correlated with neutralization potencies for polyclonal antisera from RBD-nanoparticle immunized animals (Cohen et al., 2021), this is not always true for mAbs; e.g., mAbs such as CR3022 bind to SARS-CoV-2 RBD, but neutralize only weakly or not at all (Niu et al., 2021). One mechanism by which Omicron or other RBD substitutions could indirectly affect neutralization potencies of mAbs without affecting binding to isolated RBDs is by changing the dynamics of the conversion between ‘up’ and ‘down’ conformations of RBDs on spike trimers. Some classes of anti-RBD mAbs have a strong or absolute preference for binding an ‘up’ versus a ‘down’ RBD; for example, most class 1 and class 4 anti-RBD mAbs only recognize ‘up’ RBDs (Barnes et al., 2020a). To address the ability of the mAbs investigated here to potentially recognize both ‘up’ and ‘down’ RBDs, we evaluated the accessibility of their epitopes on a spike trimer by mapping each binding epitope onto an unliganded trimer structure with one ‘up’ and two ‘down’ RBDs (PDB 6VYB) (Figure S16) and a trimer with all ‘up’ RBDs (PDB 7RKV) (Figure S17). The class 4 and 1/4 epitopes of M8a-3, M8a-6, M8a-31, M8a-34, and HSW-1 were buried when RBDs adopted the ‘down’ conformation (Figure S16A-D, F), but fully exposed in the ‘up’ RBDs (Figure S17A-D, F). Although the class 4 epitope of HSW-2 was buried in ‘down’ RBD conformation (Figure S16G) and could be partially exposed in an ‘up’ RBD conformation (Figure S17G), structural alignments showed that HSW-2 cannot bind to ‘up’ or ‘down’ RBDs in the context of a spike trimer (Figure 5F,G). By contrast, the class 3 epitope of M8a-28 was exposed in both RBD conformations (Figure S16E, S17E). Likely related to these observations, only the M8a-28–bound trimer structure showed an inter-protomer RBD distance of 31 Å (Figure 6I) that is equivalent to that of an unliganded trimer (28-40 Å) (Figure 6A). The other class 4 and 1/4 mAb Fab-bound trimer structures showed larger inter-protomer RBD distances (up to ∼70 Å), corresponding to ∼ 11 to 34 Å more outward displacement of RBDs in comparison to unliganded or class 1- or ACE2-liganded spike trimer structures (Figure 6B-H) (Barnes et al., 2020a). This outward displacement of RBDs could result in spike trimer destabilization that could lead to S1 shedding (Huo et al., 2020; Jette et al., 2021; Piccoli et al., 2020; Pinto et al., 2020).

Another property of IgG antibodies that could affect their neutralization potencies relates to their ability to utilize bivalency. Since IgG antibodies have two identical Fab arms, they can increase their apparent affinities for binding to tethered viral antigens through avidity effects, which can occur through either inter-spike crosslinking (simultaneous binding of two neighboring spike trimers) or intra-spike crosslinking (simultaneous binding of two neighboring RBDs within the same spike trimer). Inter-spike crosslinking is likely possible for most anti-RBD antibodies based on the numbers and densities of SARS-CoV-2 spikes (Barnes et al., 2020b), and intra-spike crosslinking could also occur for some anti-RBD antibodies, depending on their epitopes and binding poses (Barnes et al., 2020a). To evaluate whether the M8a or HSW mAbs could enhance their binding through intra-spike crosslinking, we measured distances between neighboring Fabs in the Fab-spike structures to predict if simultaneous binding of two Fabs within an intact IgG to adjacent RBDs on a trimer would be possible. We previously defined a distance of ≤65 Å between the C-termini of the C_H_1 domains of adjacent bound RBD-bound Fabs as being required to allow the N-termini of the two chains of an IgG hinge to each of the C-termini of two bound Fabs (Barnes et al., 2020a) and compared that to the distances measured in a previously characterized class 1/4 mAb, C118, in complex with the spike trimer (Figure 6B) (Jette et al., 2021). The measured distances in spike trimers complexed with the M8a-3 (126 Å, 130 Å, and 159 Å) (Figure 6C), M8a-34 (107 Å, 110 Å, and 150 Å) (Figure 6G), or M8a-28 (144 Å) (Figure 6I) Fabs were too large to permit intra-spike crosslinking. Although we could not measure analogous distances in the M8a-6–spike structure because only one Fab was bound (Figure 6D), the similar epitope and pose for M8a-3 and M8a-6 (Figure 3A,B; 4B,C) suggest that an IgG version of M8a-6 would also be unable to crosslink adjacent RBDs. Thus, the weak binding of M8a-6 to a spike trimer could not be improved by intra-spike crosslinking avidity effects, again rationalizing its lack of neutralizing activity (Figure 2B). For spike trimers in complex with M8a-31 Fab (Figure 6E,F), the distances between the C-termini of adjacent C_H_1 domains were measured as 52 Å and 49 Å for M8a-31 Fab bound to the WA1 and Omicron BA.1 spikes, respectively, suggestive of potential intra-spike crosslinking. We could not evaluate the potential for intra-spike crosslinking for the HSW-1 or HSW-2 mAbs because either only one Fab was bound per spike (HSW-1) (Figure 5A) or the reconstructions showed Fab binding to dissociated S1 monomer only (HSW-2) (Figure 5C).

We also used modeling to assess how the RBD-nanoparticles used to elicit the mAbs investigated here might engage with bivalent B cell receptors. To address this issue, we asked whether the geometric arrangement of RBDs on mosaic-8 RBD-mi3 nanoparticles would permit bivalent engagement of neighboring RBDs by IgGs, here representing membrane-bound B cell receptors hypothesized to engage adjacent RBDs (Figure 1D). We first constructed IgG models of each of the Fabs in the M8a and HSW Fab-spike structures (Figure 3,5). Next, we asked if it was sterically possible for both Fabs of an IgG to interact with the epitope identified from its cryo-EM structure on adjacent RBDs on a modeled RBD-mi3 nanoparticle. For each of the seven mAb epitopes, we found that the RBD-mi3 nanoparticle geometry was predicted to allow simultaneous recognition of adjacent RBDs by both Fabs of an IgG (Figure S18), thus confirming that the geometric arrangement of RBD attachment sites on SpyCatcher-mi3 would allow B cell receptor engagement through avidity effects (Figure 1D).

### mAbs elicited by mosaic-8 RBD-nanoparticles resemble EUA-approved therapeutics or a potent cross-reactive human class 1/4 anti-RBD antibody

Although our B cell cloning study from mosaic-8 RBD-nanoparticle immunized mice started with <10,000 cells, the five M8a mAbs identified as binding two or more RBDs during screening all target the desired more conserved epitopes (class 3 and class 1/4) rather than the class 1 and class 2 RBD epitopes more commonly elicited by vaccination or infection (Greaney et al., 2021a; Greaney et al., 2021b; Piccoli et al., 2020) (Figure 3,4), and three of the five mAbs were potently and/or broadly neutralizing (Figure 2B). By contrast, the only two mAbs isolated from homotypic SARS-CoV-2 RBD-nanoparticle immunized mice that were identified as binding two or more RBDs during screening targeted different epitopes (Figure 5) and were only weakly- or non-neutralizing (Figure 2B). This is consistent with our previous results showing that mosaic-8 RBD-nanoparticles are more likely to elicit cross-reactive and cross-neutralizing anti-sarbecovirus antibodies against conserved epitopes than homotypic RBD-nanoparticles (Cohen et al., 2022) and argues against use of homotypic RBD-nanoparticles as immunogens to elicit cross-reactive neutralizing antibodies.

Human mAbs that received Emergency Use Authorization for COVID-19 treatment include class 1 and class 2 anti-RBD mAbs that are no longer effective against SARS-CoV-2 variants, and class 3 anti-RBD mAbs, two of which, Bebtelovimab and Cilgavimab, retain at least partial efficacy against Omicron variants (Figure 7A,B). The class 3 RBD epitope, although less affected by SARS-CoV-2 VOC substitutions than the immunodominant class 1 and class 2 RBD epitopes, nevertheless encompasses an RBD region that includes variability across sarbecovirus sequences (Figure 1A,B, 7C-G). The ability of the class 3 epitope to accumulate substitutions is likely because the class 3 RBD region is accessible on both ‘up’ and ‘down’ RBD conformations, thus not being constrained to avoid substitutions because of contacts with other regions of a sarbecovirus spike trimer (Figure 7A) (Cohen et al., 2022). The epitope identified for M8a-28 (Figure 7C) resembles epitopes of the class 3 anti-RBD therapeutic mAbs (Figure 7D-G). Some of these mAbs, including M8a-28 (Figure 2B), neutralize Omicron VOCs, but their epitope locations within a region that varies among sarbecoviruses suggests that future SARS-CoV-2 variants are likely to include substitutions that reduce or completely abrogate their efficacies (Figure 7C-G). By contrast, the more occluded class 1/4 RBD epitope (Figure 7A), to which bound mAbs can inhibit ACE2 binding (Jette et al., 2021; Liu et al., 2020a; Tortorici et al., 2021), exhibits less variability across sarbecoviruses, likely because substitutions that affect its contacts as a ‘down’ RBD with other spike trimer regions limit its variability between SARS-CoV-2 VOCs and other sarbecoviruses (Cohen et al., 2022). The fact that four of five mouse mAbs identified as binding to two or more different RBDs during B cell screening after mosaic-8 immunization target the class 1/4 epitope, in common with the potent, cross-reactive, and protective S2X259 human mAb (Tortorici et al., 2021) (Figure 7H), supports the potential for using mosaic RBD-nanoparticles as immunogens to efficiently elicit cross-reactive and potent neutralizing mAbs against SARS-CoV-2 variants and animal sarbecoviruses that could spill over to infect humans. In addition, together with previous challenge and serum epitope mapping (Cohen et al., 2022), these results further validate mosaic-8 RBD-nanoparticles as a broadly protective vaccine candidate.

## Discussion

Here, we characterized mouse mAbs elicited using mosaic RBD-nanoparticles (M8a mAbs) or homotypic RBD-nanoparticles (HSW mAbs) using both structural and functional analyses, showing that mosaic RBD-nanoparticles induce potently neutralizing antibodies that cross-react between animal sarbecoviruses and SARS-CoV-2 VOCs (Figure 2). Although we identified only five mAbs that bound to two or more RBDs from mosaic-8 nanoparticle immunized mice in these first experiments, one mAb (M8a-3) was both cross-reactive and strongly neutralizing, and two others (M8a-31 and M8a-34) were less potently neutralizing but were cross-reactive (Figure 2), demonstrating the potential for the mosaic-8 immunogen to efficiently elicit potentially useful therapeutic antibodies that could be identified in larger-scale experiments and developed as future therapeutic mAbs. Another mAb (M8a-28) potently neutralized SARS-CoV-2 variants and resembled therapeutic antibodies in current use (Figure 7C-G).

Structural studies of Fab complexes with SARS-CoV-2 spike trimers, including one with Omicron BA.1, demonstrated that four of the five mAbs isolated from mosaic-8 immunized mice recognized conserved epitopes (Figure 4), as designed in the immunization approach (Figure 1D) and as shown for polyclonal antisera raised in mice by mosaic-8 RBD-nanoparticle immunization (Cohen et al., 2022). By contrast, antibodies raised in homotypic SARS-CoV-2 RBD-nanoparticle immunized mice more commonly recognize variable class 1 and class 2 RBD epitopes (Cohen et al., 2022), likely explaining why it was more difficult in the current study to isolate single B cells from homotypic RBD-nanoparticle immunized mice secreting IgGs that bound two or more labeled RBDs (Figure 2A, Figure S3). The two cross-RBD binding mAbs we were able to isolate from homotypic RBD-nanoparticle immunized mice showed binding to multiple sarbecovirus RBDs (Figure 2A) but were only weakly- or non-neutralizing (Figure 2B). Structural studies rationalized the weak neutralization profiles of the HSW mAbs: the HSW-1–spike structure showed only one bound Fab per trimer (Figure 5A) as compared with three bound Fabs per trimer in the structures of more potently neutralizing mAbs (Figure 3), and the HSW-2 Fab epitope (Figure 5D) was incompatible with binding to its RBD epitope on intact spike trimer (Figure 5F,G), resulting in a trimer dissociation and a Fab–monomeric S1 subunit structure (Figure 5C).

Together with previous studies (Cohen et al., 2021; Cohen et al., 2022), the current results support the mosaic-8 RBD-nanoparticles as a promising vaccine candidate to protect against SARS-CoV-2, including future variants, and against potential spillover sarbecoviruses from animal reservoirs. In addition, the fact that potent cross-reactive mAbs were identified from relatively few B cells suggest that high-throughput screening of larger samples from animals immunized with mosaic-8 RBD-mi3 could be used to identify many new therapeutic mAbs, which could then be used to prevent or treat infections of Omicron and future SARS-CoV-2 variants.

## Materials and methods

### Preparation of homotypic and mosaic-8 RBD-mi3 nanoparticles

Mammalian expression vectors encoding RBDs of SARS-CoV-2 and other sarbecoviruses were constructed as described (Cohen et al., 2021) in two versions: one with a C-terminal 6x-His tag and a SpyTag003 (RGVPHIVMVDAYKRYK) (Keeble et al., 2019) for the 8 RBDs that were coupled to SpyCatcher003-mi3 nanoparticles (Keeble et al., 2019) and other versions with only a 6x-His tag or with a His tag plus an Avi tag for ELISAs. Expression vectors encoding RBDs were constructed similarly for the following sarbecoviruses: BM4831-CoV (GenBank NC014470; spike residues 310-530), BtKY72-CoV (GenBank KY352407; spike residues 309-530), C028 (GenBank AAV98001.1; spike residues 306-523), Khosta2 (GenBank QVN46569.1; spike residues 307-526), LYRa3 (GenBank AHX37569.1; spike residues 310-527), Pangolin17-CoV (GenBank QIA48632; spike residues 317-539), RaTG13-CoV (GenBank QHR63300; spike residues 319-541), Rf1-CoV (GenBank DQ412042; spike residues 310-515), RmYN02-CoV (GISAID EPI_ISL_412977; spike residues 298-503), Rs4081-CoV (GenBank KY417143; spike residues 310-515), RshSTT200 (GISAID EPI_ISL_852605; spike residues 306-519), SARS-CoV (GenBank AAP13441.1; spike residues 318-510), SARS-CoV-2 WA1 (GenBank MT246667.1; spike residues 319-539), SHC014-CoV (GenBank KC881005; spike residues 307-524), W1V1-CoV (GenBank KF367457; spike residues 307-528), and Yun11-CoV (GenBank JX993988; spike residues 310-515). SARS-CoV-2 variants, including Beta, Delta, Omicron BA.1, and Omicron BA.2 with C-terminal 6x-His tag were also constructed similarly as the SARS-CoV-2 WA1 RBD construct for ELISA. All RBD proteins were expressed by transient transfection of Expi293F cells and purified by Ni-NTA and size exclusion chromatography (SEC) using a HiLoad 16/600 Superdex 200 column (Cytiva, Marlborough, MA) (Barnes et al., 2020b). Peak fractions were pooled, concentrated, and stored at 4°C until use.

SpyCatcher003-mi3 (Keeble et al., 2019) were expressed in *E. coli* BL21 (DE3)-RIPL (Agilent Technology) and purified as described previously (Cohen et al., 2021). Briefly, *E. coli* transduced with a SpyCatcher003-mi3 expression plasmid (Addgene, Watertown, MA) were lysed with a cell disrupter in the presence of 2 mM PMSF. After spinning at 21,000 x *g* for 30 minutes, supernatant containing SpyCatcher003-mi3 particles was passed over a pre-packed Ni-NTA column. The eluent was concentrated and further purified by SEC using a HiLoad 16/600 Superdex 200 column (Cytiva, Marlborough, MA). Peak fractions were pooled and stored at 4°C until use. SpyCatcher003-mi3 particles were used for SpyTagged RBD conjugation for up to a month after clarification by filtering using a 0.2 µm filter or spinning at 21,000 x g for 10 min.

For conjugation, purified SpyCatcher003-mi3 was incubated with purified SpyTagged RBDs (either 8 different RBDs to make mosaic-8 RBD-mi3 or SARS-CoV-2 RBD only to make homotypic RBD-mi3) at a molar ratio of 1:1.2 overnight at room temperature. Conjugation efficiencies of individual RBDs to SpyCatcher003-mi3 were verified as shown in Figure S2 of (Cohen et al., 2021). Conjugated mi3-RBD particles were purified by SEC using a Superose 6 10/300 column (Cytiva, Marlborough, MA). Peak fractions pooled and the concentrations of conjugated mi3 particles were determined using a Bio-Rad Protein Assay (Bio-Rad, Hercules, CA). Conjugated nanoparticles were characterized by electron microscopy imaging and SEC as shown in Figure S1C-E, and by electron microscopy, SEC and dynamic light scattering as shown in Figure S1 of (Cohen et al., 2022).

For negative-stain electron microscopy imaging of mosaic-8 and homotypic SARS-CoV-2 RBD-nanoparticles: ultrathin, holey carbon-coated, 400 mesh Cu grids (Ted Pella, Inc.) were glow discharged (60 s at 15 mA), and a 3 μL aliquot of SEC-purified RBD-nanoparticles was diluted to ∼40-100 µg/mL and applied to grids for 60 s. Grids were negatively stained with 2% (w/v) uranyl acetate for 30 s, and images were collected with a 120 keV FEI Tecnai T12 transmission electron microscope at 42,000x magnification.

### Immunizations

Immunizations were done using protocols, #19023, approved by the City of Hope IACUC committee. Experiments were conducted using 4–6-week-old female C57BL/6 mice (Charles River Laboratories, Wilmington, MA). Immunizations were carried out as previously described (Cohen et al., 2021) using intraperitoneal injections of 5 µg of conjugated RBD-mi3 nanoparticle (calculated as the mass of the RBD, assuming 100% efficiency of conjugation to SpyCatcher003-mi3) in 100 µL of 50% v/v AddaVax^TM^ adjuvant (Invivogen, San Diego, CA). Animals were boosted 4 weeks after the prime with the same quantity of antigen in adjuvant. A final booster was administered intraperitoneally 3 days before mouse spleen harvest.

### Beacon

Plasma B cells were isolated from immunized animals for characterization on a Berkeley Lights Beacon instrument. Spleens were isolated from two immunized mice per condition and prepared into single cell suspensions as described (Cohen et al., 2021). Plasma B cells were isolated by CD138^+^ cell enrichment (Miltenyi Biotec CD138^+^ plasma cell isolation kit, catalog no. 130-092-530). Enriched plasma B cell samples were loaded onto an OptoSelect 11k chip (Berkeley Lights, Inc., Emeryville, CA) in BLI Mouse Plasma Cell Media (Berkeley Lights, Inc., catalog no. 750-70004). Single cells were then isolated in individual nanoliter-volume compartments (Nanopens using light-based OptoElectro Positioning (OEP) manipulation with settings optimized for plasma B cells. From Mosaic-8 RBD-nanoparticle immunized animals, 9,695 cells were penned in one chip, of which 7,747 were single cell pens. For homotypic SARS-CoV-2 RBD-nanoparticle immunized animals, 9,130 cells were penned in a second chip, of which 7699 were single cell pens (Figure S3A). On chip fluorescence assays were used to identify cells secreting antibodies specific to RBD antigens. Briefly, C-terminally Avi-tagged RBDs were modified with site-specific biotinylation (Avidity, LLC, Aurora, CO) according to the manufacturer’s protocol and immobilized on streptavidin-coated beads (Berkeley Lights, Inc. catalog no. 520-00053). Assays were conducted by mixing beads coupled with one of four RBDs used for screening with a fluorescently labeled goat anti-mouse secondary antibody Alexa568 at 1:2500 dilution and importing this assay mixture into the OptoSelect 11k chip. Assays were conducted post 30 minutes incubation after cell penning at 36 °C. Images were acquired every 5 minutes for 9 cycles while the beads remained stationary in the main channel above the Nanopens of the OptoSelect chip. Antibodies specific for the immobilized RBD bound the antigen-coupled beads, which sequestered the fluorescent secondary antibody, creating a “bloom” of fluorescent signal immediately above Nanopens containing plasma B cells. Beads were washed out of the chip, and this assay was conducted for each of the four RBDs. After completion of all assays, RBD-specific cells of interest were exported using OEP from individual nanopen chambers to individual wells of a 96-well PCR plate containing lysis buffer.

After running assays and selecting positive blooms with single cells, we ran the OptoSeq BCR Export workflow, which performs reverse transcription overnight on the chip and exports cell lysates containing cDNA on capture beads onto a 96 well plate. cDNA amplification and chain-specific PCR were performed the following day and run on an agarose gel to confirm that bands of the correct size were present. PCR products were then purified using AMPure XP magnetic beads and submitted for Sanger sequencing at the City of Hope Sequencing Core.

### Cloning

Sequences for V_H_ and V_L_ domains were codon optimized using GeneArt (Thermo Fisher Scientific, Waltham, MA) and gene blocks for each domain were purchased from Integrated DNA Technologies (IDT, Coralville, IA). Expression constructs were assembled using Gibson reactions (Gibson et al., 2010; Gibson et al., 2009). The heavy chain for IgG expression was constructed by subcloning the V_H_ gene into a p3BNC expression vector encoding the human IgG C_H_1, C_H_2, and C_H_3 domains, and the heavy chain for Fab expression was constructed by assembling the V_H_ gene into a p3BNC expression vector encoding a human C_H_1 and a C-terminal 6x-His tag. The expression plasmid for the light chain was constructed by subcloning the V_L_ gene into a p3BNC vector that also encoded kappa human C_L_. The numbering of V_H_ and V_L_ protein sequences and the identification of the V gene segments were determined using the ANARCI server (Dunbar et al., 2016).

### IgG and spike trimer production and purification

Proteins were expressed in Expi293 cells by transient transfection. IgGs and a previously-described human ACE2-Fc construct (Jette et al., 2021) were purified from cell supernatants using MabSelect SURE columns (Cytiva, Marlborough, MA), and His-tagged Fabs were isolated from cell supernatants using Ni-NTA columns (Qiagen, Hilden, Germany). IgGs, ACE2-Fc, and Fabs were further purified by SEC using a Superdex 200 16/600 column (Cytiva, Marlborough, MA). Purified proteins were concentrated using a 100 kDa and 30 kDa cutoff concentrator (EMD Millipore, Burlington, MA), respectively, to 10 to 15 mg/mL, and final concentrated proteins were stored at 4 °C until use. 6P versions (Hsieh et al., 2020) of soluble SARS-CoV-2 WA1 and SARS-CoV-2 Omicron BA.1 spike trimers were isolated from cell supernatants using a Ni-NTA column (Qiagen, Hilden, Germany). Eluents from Ni-NTA purifications were subjected to SEC using a HiLoad Superdex 200 16/600 column followed by a Superose 6 10/300 (Cytiva, Marlborough, MA) column. Peak fractions were pooled and concentrated to ∼6 mg/ml, flash frozen in 50 µL aliquots, and stored at −80 °C until use.

### ELISAs

Nunc® MaxiSorp™ 384-well plates (Millipore Sigma, St. Louis, MO) were coated with 10 µg/mL of purified RBD in 0.1 M NaHCO_3_ pH 9.8 and stored overnight at 4 °C. After blocking with 3% bovine serum albumin (BSA) for an hour at room temperature, plates were washed with Tris-buffered saline including 0.1% Tween 20 (TBST). After removing blocking solution from the plates, 100 μg/mL of purified IgGs were serially diluted by 4-fold using TBST with 3% BSA and incubated with plates at room temperature for 3 hours. Plates were then washed with TBST and incubated with secondary HRP-conjugated goat anti-human IgG (Southern Biotech, Birmingham, AL) at a 1:15,000 dilution for 45 minutes at room temperature. Plates were washed with TBST, developed using SuperSignal ELISA Femto Maximum Sensitivity Substrate (Thermo Fisher Scientific, Waltham, MA), and read at 425 nm. ELISA data were collected in duplicate, and each assay was conducted at least twice for the seven mAbs that were structurally characterized. Curves were plotted and integrated to obtain half-maximal effective concentrations (EC_50_) using Graphpad Prism v9.3.1.

Competition ELISAs were performed using a Tecan Evo liquid handling robot using modifications of a previously described protocol (Escolano et al., 2021). IgGs were randomly biotinylated at primary amines using EZ-link NHS-PEG4 Biotinylation Kit according to the manufacturer’s protocol (Thermo Fisher Scientific, Waltham, MA). SARS-CoV-2 RBD (2.5 µg/mL) was adsorbed overnight at 4°C to a 384-well Nunc MaxiSorp ELISA plate (Millipore Sigma, St. Louis, MO). The RBD was removed via aspiration and the plate blocked with 3% BSA in TBST for 1 hour at room temperature. The blocking was removed via aspiration and 10 µg/mL unlabeled IgG was added and incubated for 2 hours, followed by addition of 0.25 µg/mL biotinylated IgG. The plate was incubated for 2 hours at room temperature, and bound biotinylated IgG was detected using horseradish peroxidase-conjugated streptavidin (Southern Biotech, Birmingham, AL) (1 hour, room temperature) and developed with SuperSignal ELISA Femto Substrate (Thermo Fisher Scientific, Waltham, MA). Relative light units (RLU) were measured and the signal for each competition pair was normalized to the signal for the biotinylated IgG when unlabeled IgG was not present. Measurements were performed in technical quadruplicates. Data presented are representative of two independent experiments.

### Neutralization assays

SARS-CoV-2, SARS-CoV-2 VOCs, SARS-CoV, WIV1, SHC014, BtKY72 (including mutations allowing human ACE2 binding (Starr et al., 2022)), Khosta2/SARS-CoV Chimera, and LyRa3/SARS-CoV Chimera pseudoviruses based on HIV lentiviral particles were prepared as described (Crawford et al., 2020; Robbiani et al., 2020). Khosta2/SARS-CoV and LyRa3/SARS-CoV Chimeras were constructed by replacing the RBD of SARS-CoV Spike with the RBDs of Khosta2 and LyRa3 Spikes separately. Assays were done using 4-fold dilutions of purified IgGs at a starting concentration of 100 μg/mL by incubating with a pseudovirus at 37 °C for an hour. After incubating with 293TACE2 target cells for 48 hours at 37 °C, cells were washed 2 times with phosphate-buffered saline (PBS) and lysed with Luciferase Cell Culture Lysis 5x reagent (Promega, Madison, WI). Using the Nano-Glo Luciferase Assay System (Promega, Madison, WI), the NanoLuc Luciferase activity in lysates was measured. Relative luminescence units (RLUs) were normalized to values derived from cells infected with pseudovirus in the absence of IgG. Data were collected at each IgG concentration in duplicate and reported data come from assays performed at least twice (except for the M8a-28 against Omicron BA.1 assay, which was performed once). Half-maximal inhibitory concentrations (IC_50_ values) were determined using nonlinear regression in AntibodyDatabase (West et al., 2013).

### X-ray crystallography

RBD-Fab complexes were formed by incubating SARS-CoV-2 RBD with a 1.1x molar excess of Fab for an hour at room temperature. Complexes were purified by SEC using a Superdex 200 10/300 Increase column (Cytiva, Marlborough, MA). Peak fractions containing RBD-Fab complexes were pooled and concentrated to ∼15 mg/ml. Crystallization trials were set up using commercially available screens by mixing 0.2 μL of RBD-Fab complex and 0.2 μL well solution using a TTP LabTech Mosquito instrument via the sitting drop vapor diffusion method at room temperature. Crystals of M8a-6 Fab–RBD complex were obtained from Proplex screen (Molecular Dimensions, Holland, OH), containing 0.1 M sodium citrate pH 5.5 and 15 % PEG 6,000. Crystals of M8a-34 Fab–RBD complex were obtained from a PEGion screen (Hampton Research, Aliso Viejo, CA), containing 2% v/v tacsimate pH 4.0, 0.1 M sodium acetate trihydrate pH 4.6, 16 % PEG 3,350. Crystals of RBD–HSW-2 complexes were obtained from a Proplex screen (Molecular Dimensions, Holland, OH), containing 0.2 M sodium chloride, 0.1 M sodium/potassium phosphate pH 6.5, 25 % PEG 1,000. All crystals were cryoprotected in well solution mixed with 20% glycerol or PEG 400 before freezing in liquid nitrogen.

X-ray diffraction data were collected at the Stanford Synchrotron Radiation Lightsource (SSRL) beamline 12-2 with a Pilatus 6M pixel detector (Dectris, Philadelphia, PA) using the Blu-ice interface (McPhillips et al., 2002) (Table S3). All X-ray datasets were indexed and integrated with XDS (Kabsch, 2010) and scaled with Aimless (Winn et al., 2011). The M8a-6 Fab–RBD structure was solved by molecular replacement using a structure of a Fab-RBD complex from a single-particle cryo-EM structure (PDB 7SC1) as the input model for *Phaser* in Phenix (Liebschner et al., 2019). During the refinement of the M8a-6 Fab–RBD structure, we observed electron density for a second RBD and the variable domains of M8a-6 Fab, but no Fab constant domains were found. Refinement of a model containing the original M8a-6 Fab–RBD complex, a second copy of RBD and the variable domains resulted in no improvements in the refinement statistics. We thus only partially refined the coordinates for the M8a-6 Fab–RBD crystal structure, which were then docked and refined in the cryo-EM M8a-6–spike reconstruction. The M8a-34 Fab–RBD structure was solved by molecular replacement using the partially refined model of M8a-6–RBD complex structure as the input model for *Phaser* in Phenix (Liebschner et al., 2019). The HSW-2 Fab–RBD structure was solved by molecular replacement using the partially refined model of M8a-34–RBD complex structure as the input model for *Phaser* in Phenix (Liebschner et al., 2019). Iterative refinement and model-building cycles were carried out with phenix.refine in Phenix (Liebschner et al., 2019) and *Coot* (Emsley et al., 2010), respectively. The refined models were subsequently used as input models for docking into cryo-EM maps of Fab-spike complexes.

### Cryo-EM Sample Preparation

SARS-CoV-2 S–Fab complexes were formed by incubating purified spike trimer and Fabs at a 1.1x molar excess of Fab per spike protomer at room temperature for 30 minutes to a final concentration of ∼2 mg/mL. Fluorinated octylmaltoside solution (Anatrace, Maumee, OH) was added to the spike-Fab complex to a final concentration of 0.02% (w/v) prior to freezing, and 3 µL of the complex/detergent mixture was immediately applied to QuantiFoil 300 mesh 1.2/1.3 grids (Electron Microscopy Sciences, Hatfield, PA) that had been freshly glow discharged with PELCO easiGLOW (Ted Pella, Redding, CA) for 1 min at 20 mA. Grids were blotted for 3 to 4 seconds with 0 blot force using Whatman No.1 filter paper and 100% humidity at room temperature and vitrified in 100% liquid ethane using a Mark IV Vitrobot (Thermo Fisher Scientific, Waltham, MA).

### Cryo-EM data collection and processing

Single-particle cryo-EM datasets for complexes of SARS-CoV-2 WA1 spike 6P with M8a-3 Fab, M8a-6 Fab, M8a-28 Fab, M8a-31 Fab, M8a-34 Fab or HSW-1 Fab and SARS-CoV-2 Omicron BA.1 spike 6P with M8a-31 Fab were collected using SerialEM automated data collection software (Mastronarde, 2005) on a 300 keV Titan Krios (Thermo Fisher Scientific, Waltham, MA) cryo-electron microscope equipped with a K3 direct electron detector camera (Gatan, Pleasanton, CA). For SARS-CoV-2 WA1 spike 6P complexed with HSW-2, a dataset was collected with SerialEM (Mastronarde, 2005) on a 200 keV Talos Arctica cryo-electron microscope (Thermo Fisher Scientific, Waltham, MA) equipped with a K3 camera (Gatan, Pleasanton, CA). Movies were recorded with 40 frames, a defocus range of −1 to −3 µm, and a total dosage of 60 e^-^/Å^2^ using a 3×3 beam image shift pattern with 3 exposures per hole in the superresolution mode with a pixel size of 0.416 Å for the collections on the Krios and a single exposure per hole in the superresolution mode with a pixel size of 0.4345 Å for the collection on the Talos Arctica. Detailed data processing workflows for each complex structure are outlined in figs. S6-13. All datasets were motion corrected with patch motion correction using a bining factor of 2, and CTF parameters were estimated using Patch CTF in cryoSPARC v3.2 (Punjani et al., 2017). Particle picking was done with blob picker in cryoSPARC using a particle diameter of 100 to 200 Å, and movies and picked particles were inspected before extraction. Particles were extracted and classified using 2D classification in cryoSPARC (Punjani et al., 2017). After discarding ice and junk particles, the remaining particles were used for *ab initio* modeling with 4 volumes, which were futher refined with heterogenerous refinement in cryoSPARC (Punjani et al., 2017). Subsequent homogeneous and non-uniform refinements were carried out for final reconstructions in cryoSPARC (Punjani et al., 2017). Because Fab interactions with ‘up’ RBDs are generally not well resolved in Fab-spike complex structures (Pinto et al., 2020), we used masks to locally refine and improve the interfaces of Fabs bound to ‘up’ RBDs when necessary. For local refinements, masks were generated using UCSF Chimera (Pettersen et al., 2004) and refinements were carried out in cryoSPARC (Punjani et al., 2017).

### Cryo-EM Structure Modeling and Refinement

An initial model of the M8a-3 Fab–spike trimer complex was generated by docking a single-particle cryo-EM Fab-SARS-CoV-2 spike 6P complex structure (PDB 7SC1) into the cryo-EM density using UCSF Chimera (Pettersen et al., 2004). The model was refined using real space refinement in Phenix (Liebschner et al., 2019). The Fab amino acid seqence was manually corrected in *Coot* (Emsley et al., 2010). The model of the M8a-3 Fab–spike complex was subsequently used for docking and model generation for remaining Fab-spike trimer complexes. For the Fab-spike complexes that we have RBD-Fab crystal structures for (M8a-6 Fab-RBD, M8a-34 Fab–RBD and HSW-2 Fab–RBD structures), we first docked the spike trimer (PDB 7SC1) in the EM density map, manually fitted the RBDs in *Coot* (Emsley et al., 2010) and refined the spike trimer using phenix.real_space_refine (Liebschner et al., 2019). The RBD-Fab structures were then aligned to each of the RBDs in the corresponding Fab–spike complexes, and the RBD regions in the EM model were replaced by the RBDs from crystal structures upon structural alignments in *Coot* (Emsley et al., 2010). The final model containing the spike trimer and the Fabs were subsequently refined with phenix.real_space_refine (Liebschner et al., 2019). Iterative real space refinement and model building were separately carried out in Phenix (Liebschner et al., 2019) and *Coot* (Emsley et al., 2010). Single-particle cryo-EM refinement statistics are reported in Table S2.

### Structure Analyses

Structure figures were made using UCSF ChimeraX (Goddard et al., 2018; Pettersen et al., 2021). Distances were measured using PyMol (Schrödinger and DeLano, 2020). Interacting residues between a Fab and RBD were analyzed by PDBePISA (Krissinel and Henrick, 2007) using the following interaction definitions: potential H bonds were defined as a distance less than 3.9 Å between the donor and acceptor residues when H was present at the acceptor and there was an A-D-H angle between 90° and 270°; potential salt bridges and van der Waals interactions were defined between residues that were less than 4 Å (Krissinel and Henrick, 2007). Sequence alignments were done using Geneious (https://www.geneious.com/).

To evaluate the potential for intra-spike crosslinking by the two Fabs of a single IgG binding to adjacent RBDs within a single spike trimer, we measured the distances between the Cα atoms of the C-terminal residues of the C_H_1 domains of adjacent RBD-binding Fabs in the structures of mAb-spike complexes as described previously (Barnes et al., 2020a). A cut-off of no more than 65 Å was used to identify IgGs whose binding orientation could allow for both Fabs to bind simultaneously to adjacent RBDs in a single spike trimer. This cut-off was larger than the distance measured between comparable residues of C_H_1 domains in intact IgG crystal structures (42Å, PDB 1HZH; 48Å, PDB 1IGY; 52Å, PDB 1IGT) to account for potential influences of crystal packing, flexibilities in the elbow bend angle relating the V_H_-V_L_ and C_H_1-C_L_, and uncertainties in the placements of C_H_1-C_L_ domains in cryo-EM structures of the Fab-spike complexes (Barnes et al., 2020a).

## Acknowledgments

We thank J. Vielmetter, P. Hoffman, A. Rorick, K. Storm and the Caltech Beckman Institute Protein Expression Center for protein production, D. Veesler for BtKY72 neutralization advice, A. Gonzales for isolation and sequencing of positive B cells, the Antibody Discovery Engine and the Drug Discovery and Structural Biology Shared Facility at City of Hope, Songye Chen and the Caltech Cryo-EM facility for cryo-EM data collection, and Jens Kaiser, staff at Stanford Synchrotron Radiation Lightsource, and the Caltech Molecular Observatory for X-ray data collection support. Cryo-Electron microscopy was performed in the Beckman Institute Resource Center for Transmission Electron Microscopy at Caltech. Use of the Stanford Synchrotron Radiation Lightsource, SLAC National Accelerator Laboratory, is supported by the U.S. Department of Energy, Office of Science, Office of Basic Energy Sciences under Contract No. DE-AC02-76SF00515. The SSRL Structural Molecular Biology Program is supported by the DOE Office of Biological and Environmental Research, and by the National Institutes of Health, National Institute of General Medical Sciences (P30GM133894). The contents of this publication are solely the responsibility of the authors and do not necessarily represent the official views of NIGMS or NIH. These studies were funded by the National Institutes of Health (NIH) P01-AI138938-S1 (P.J.B.) and City of Hope’s Integrated Drug Development Venture supported by the National Cancer Institute of the National Institutes of Health P30 CA033572 (JCW), Bill and Melinda Gates Foundation INV-034638 INV-004949 (PJB), the Caltech Merkin Institute (PJB), and a George Mason University Fast Grant (PJB).

## Author contributions

C.F., A.A.C, J.R.K., J.C.W., and P.J.B. conceived the study and analyzed the data. C.F. performed single-particle cryo-electron microscopy, X-ray crystallography, interpreted structures and analyzed antibody sequences. A.A.C. prepared nanoparticles and performed negative stain electron microscopy. M.P. and A.F.H.H. isolated B cells and generated mAb sequences. C.F. and Y.E.L prepared and purified Fabs and IgGs. A.A.C., J.R.K., and Z.W. performed ELISAs. P.N.P.G., L.M.K., and A.A.C. performed neutralization assays. C.F., J.R.K., K.E.M. and P.J.B. wrote the paper with contributions from other authors.

## Data and materials availability

Atomic models and cryo-EM maps generated from cryo-EM studies of the M8a-3–WA1 spike 6P, M8a-6–WA1 spike 6P, M8a-28–WA1 spike 6P, M8a-31– WA1 spike 6P, M8a-31–Omicron BA.1 spike 6P, M8a-34–WA1 spike 6P, HSW-1–WA1 spike 6P, and HSW-2–WA1 spike S1 domain complexes have been deposited at the Protein Data Bank (PDB) and Electron Microscopy Data Bank (EMDB) under the following accession codes: PDB 7UZ4, 7UZ5, 7UZ6, 7UZ7, 7UZ8, 7UZ9, 7UZA, and 7UZB; EMDB EMD-26878, EMD-26879, EMD-26880, EMD-26881, EMD-26882, EMD-26883, EMD-26884, and EMD-26885. Atomic models generated from crystal structures of M8a-34–RBD and HSW-2–RBD complexes have been deposited at the PDB under accession codes 7UZC and 7UZD, respectively. Materials are available upon request to the corresponding author with a signed material transfer agreement. This work is licensed under a Creative Commons Attribution-NonCommercial 2.0 Generic (CC BY-NC 2.0) license, which permits unrestricted use, distribution, and reproduction in any medium, provided the original work is properly cited and not for commercial purposes. To view a copy of this license, visit https://creativecommons.org/licenses/by-nc/2.0/. This license does not apply to figures/photos/artwork or other content included in the article that is credited to a third party; obtain authorization from the rights holder before using such material.

---

**Figure S1.**
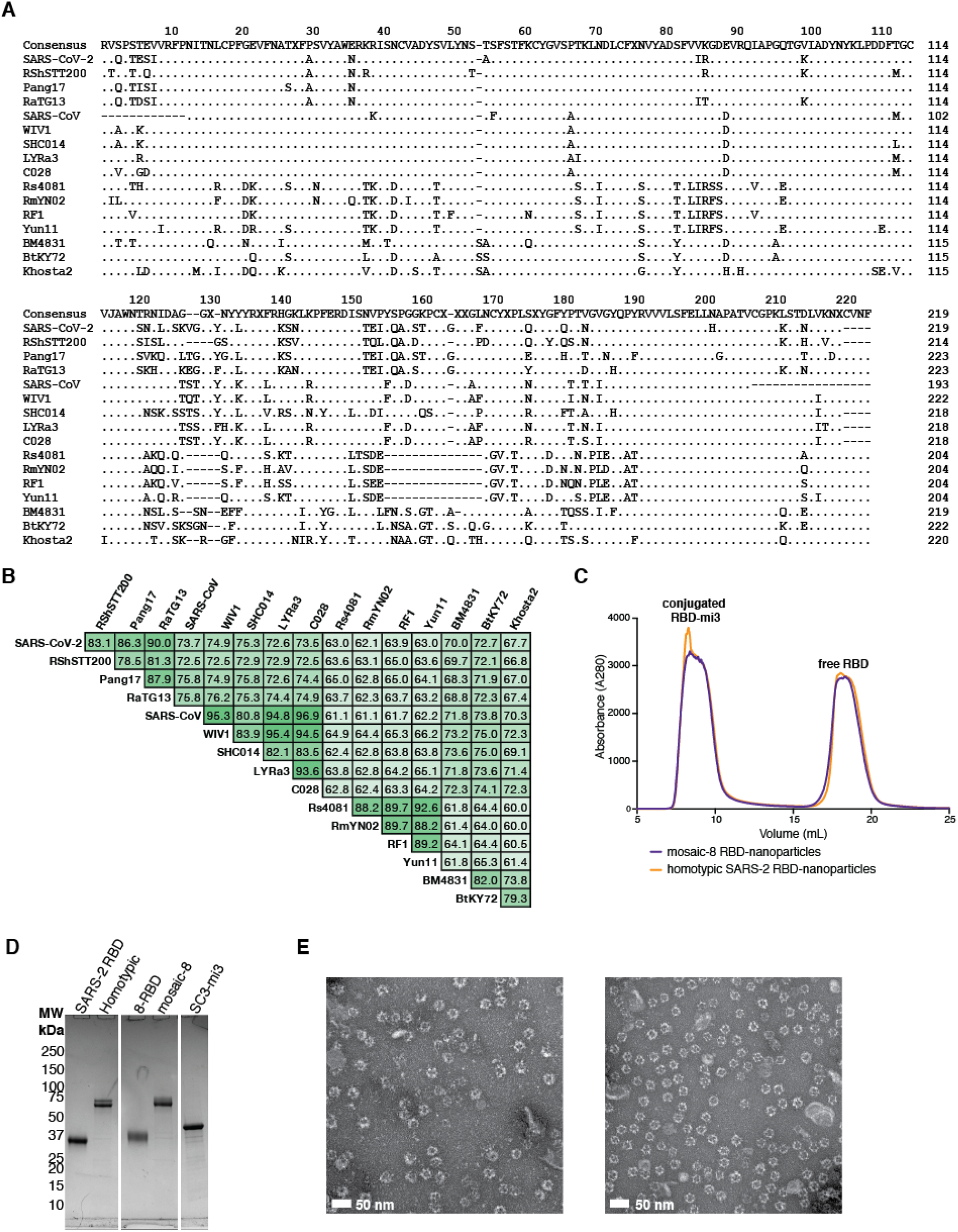
Sarbecovirus RBDs and construction of RBD-nanoparticles. (**A**) Sequence alignment of RBDs from 16 sarbecoviruses. (**B**) Pairwise sequence identities (%) relating the RBDs in panel A. (**C**) SEC profile showing separation of conjugated SpyCatcher003-mi3 nanoparticles from free RBDs. RBDs were added at a 2-fold molar excess over SpyCatcher003-mi3 subunits. (**D**) Reducing SDS-PAGE analysis of free RBD, purified conjugated RBD-nanoparticles, and unconjugated SpyCatcher-mi3 nanoparticles. (**E**) Negative stain EM of conjugated nanoparticles. Left: mosaic-8 RBD-nanoparticles. Right: homotypic SARS-CoV-2 RBD-nanoparticles.

**Figure S2.**
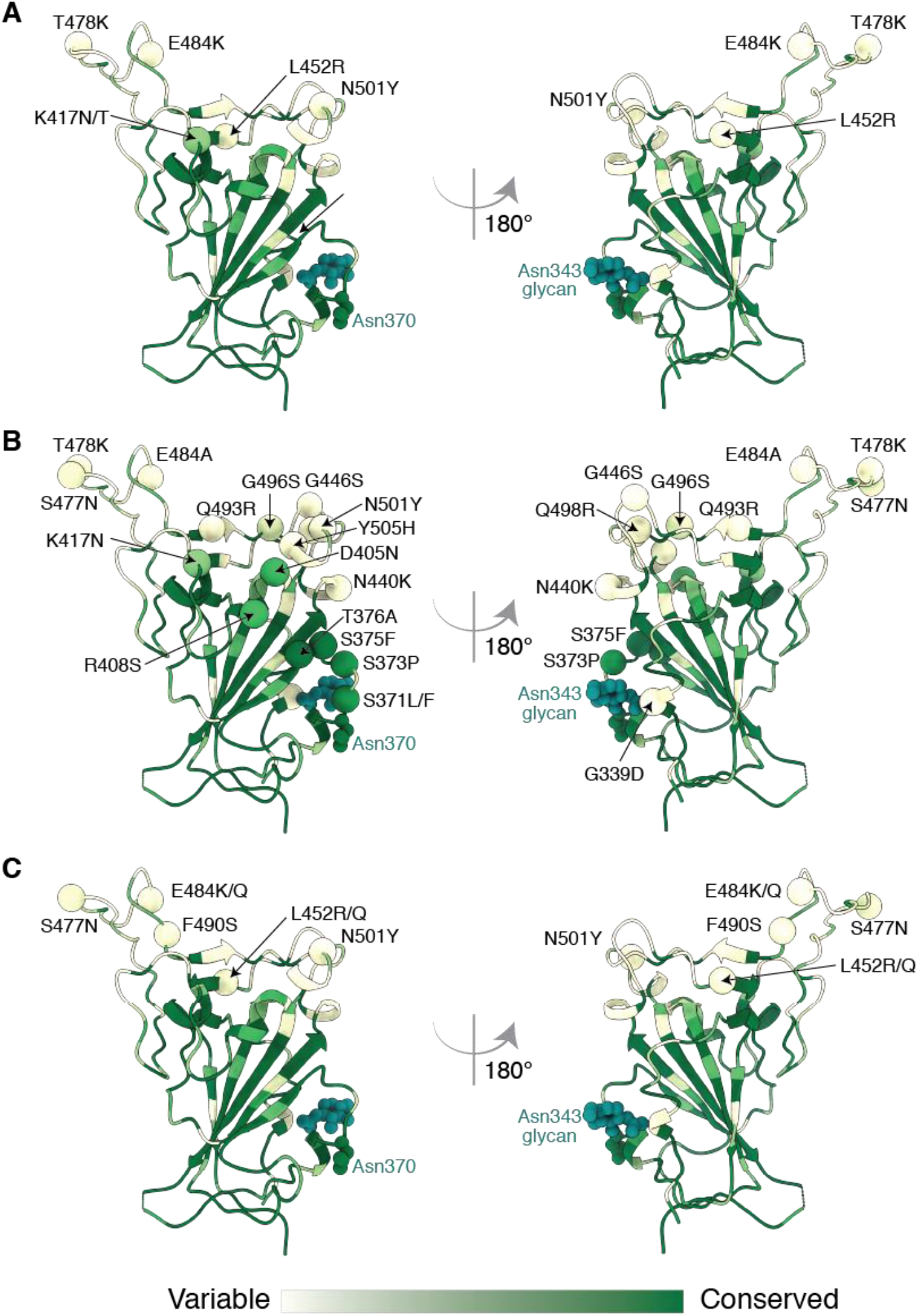
RBD VOC and VOI substitutions. Locations of RBD substitutions in VOCs and VOIs (https://viralzone.expasy.org/9556) are shown as spheres colored according to a variability gradient (bottom) on the WA1 RBD structure (PDB 7BZ5). The N-linked glycan at position 343 of SARS-CoV-2 RBD is shown as teal spheres, and a potential N-linked glycosylation site at position 370 (SARS-CoV-2 numbering) that is found in sarbecovirus RBDs but not in the SARS-CoV-2 RBD is also shown in spheres. (**A**) SARS-CoV-2 Alpha, Beta, Gamma and Delta variants. (**B**) SARS-CoV-2 Omicron BA.1 and BA.2 variants. (**C**) SARS-CoV-2 VOIs.

**Figure S3.**
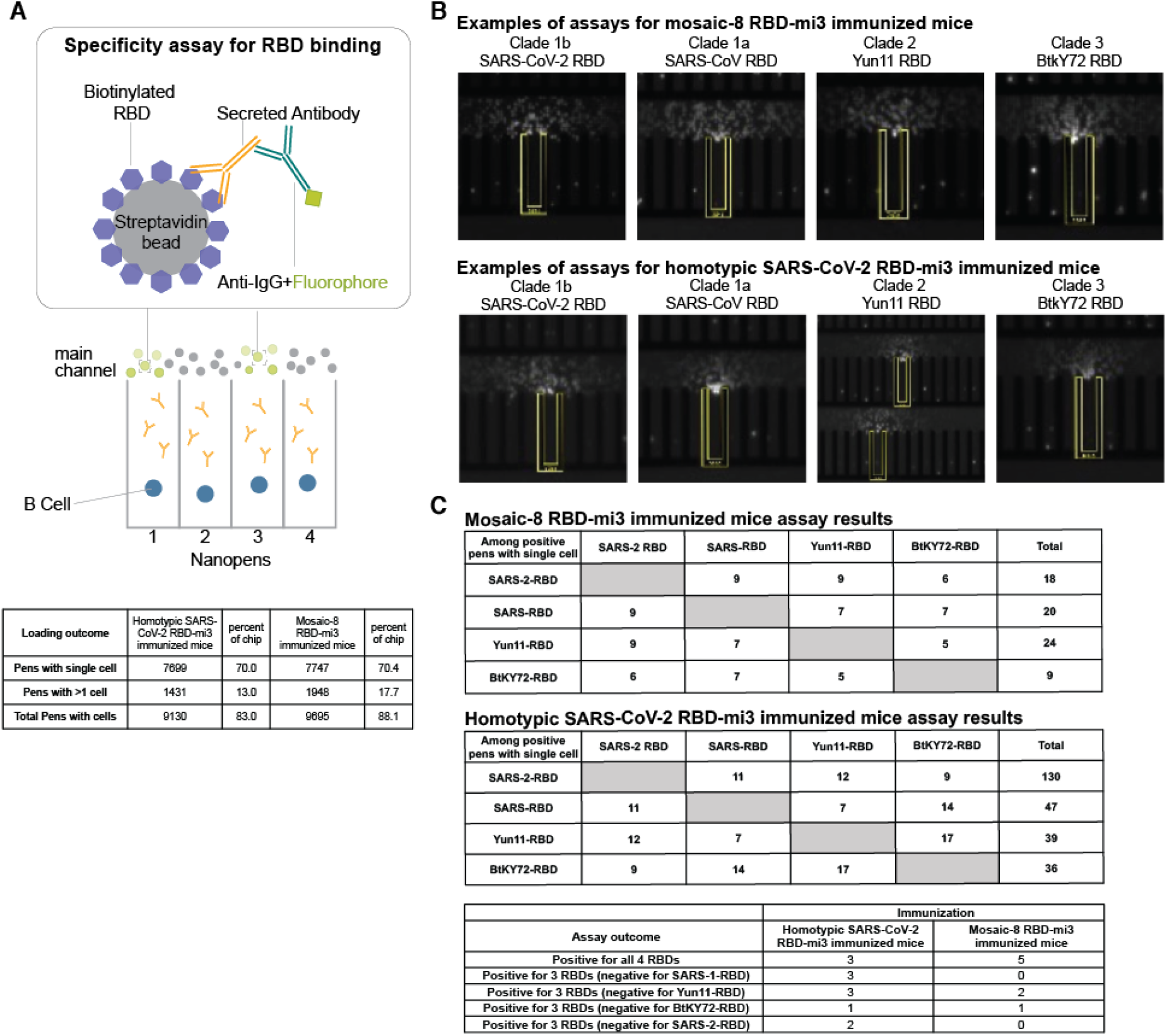
Beacon setup. Individual plasma cells were isolated, cultured, and assayed for RBD binding activities using the Beacon optofluidic system. (**A**) Antigen-specific binding activity was assayed using biotinylated RBDs immobilized on streptavidin-coated beads and loaded into the main channel of the Beacon microfluidic chip, which contained single plasma B cells in individual culture compartments (Nanopens). The specificity of secreted antibodies for the antigen presented in the channel was detected via the local concentration of a fluorescently-labeled secondary above a Nanopen secreting an antigen-specific antibody. (**B**) Examples of detection of secreted antibody binding to different RBDs. The Nanopen(s) from which the secreted antibody signal was emanating over the assay time course is indicated in yellow. (**C**) Summary of cross-reactive antigen-specificity assay results for four RBDs.

**Figure S4.**
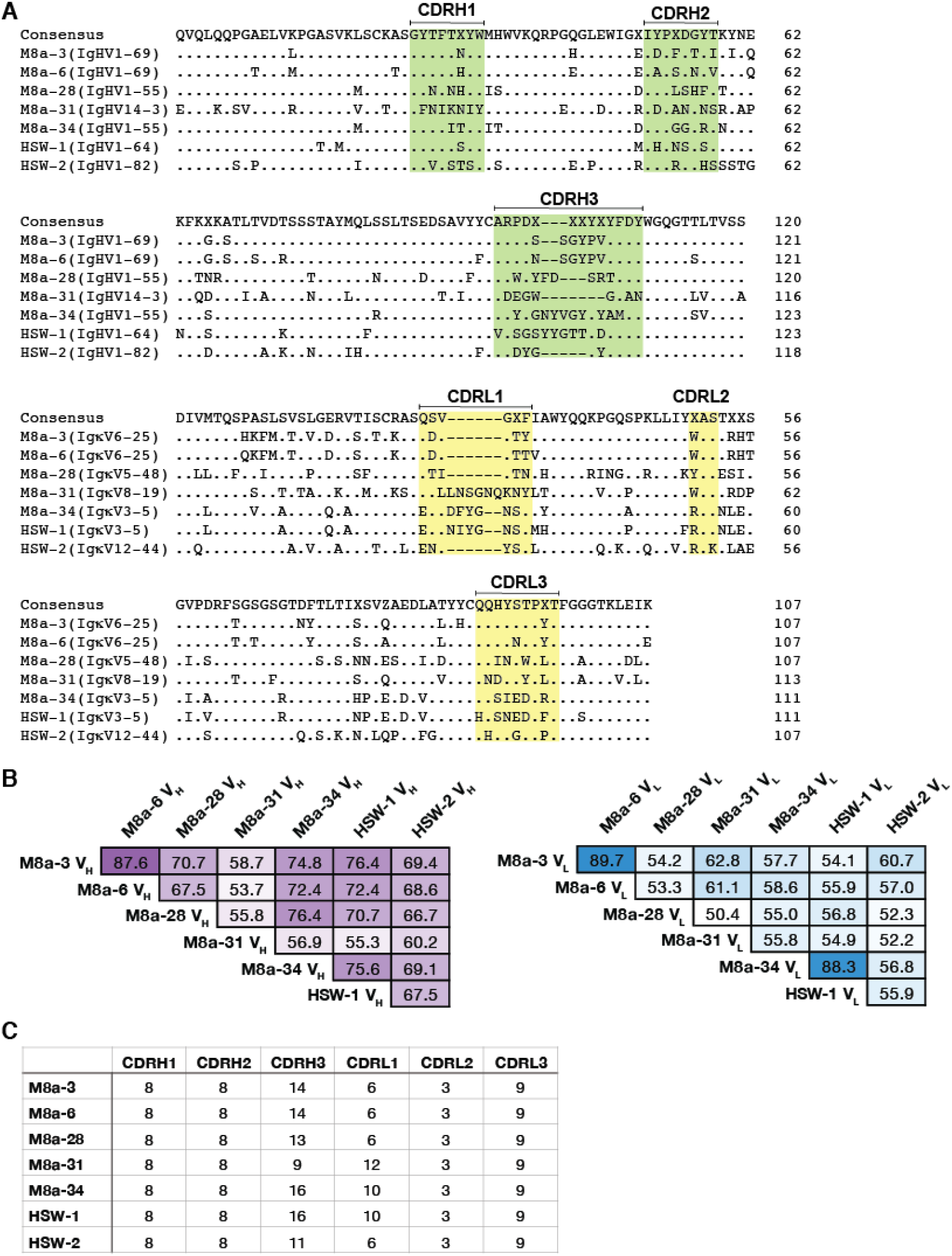
Sequence alignment of seven mAbs isolated from RBD nanoparticle-immunized mice that bound two or more sarbecovirus RBDs. (**A**) Sequence alignment of V_H_ and V_L_ domains. CDRs were assigned using IMGT definitions (Lefranc et al., 2015). VH and VL gene segment assignments for each mAb were shown in parentheses. (**B**) Pairwise sequence identities (%) between V_H_ and V_L_ domains in (**A**). (**C**) Number of residues in CDRs.

**Figure S5.**
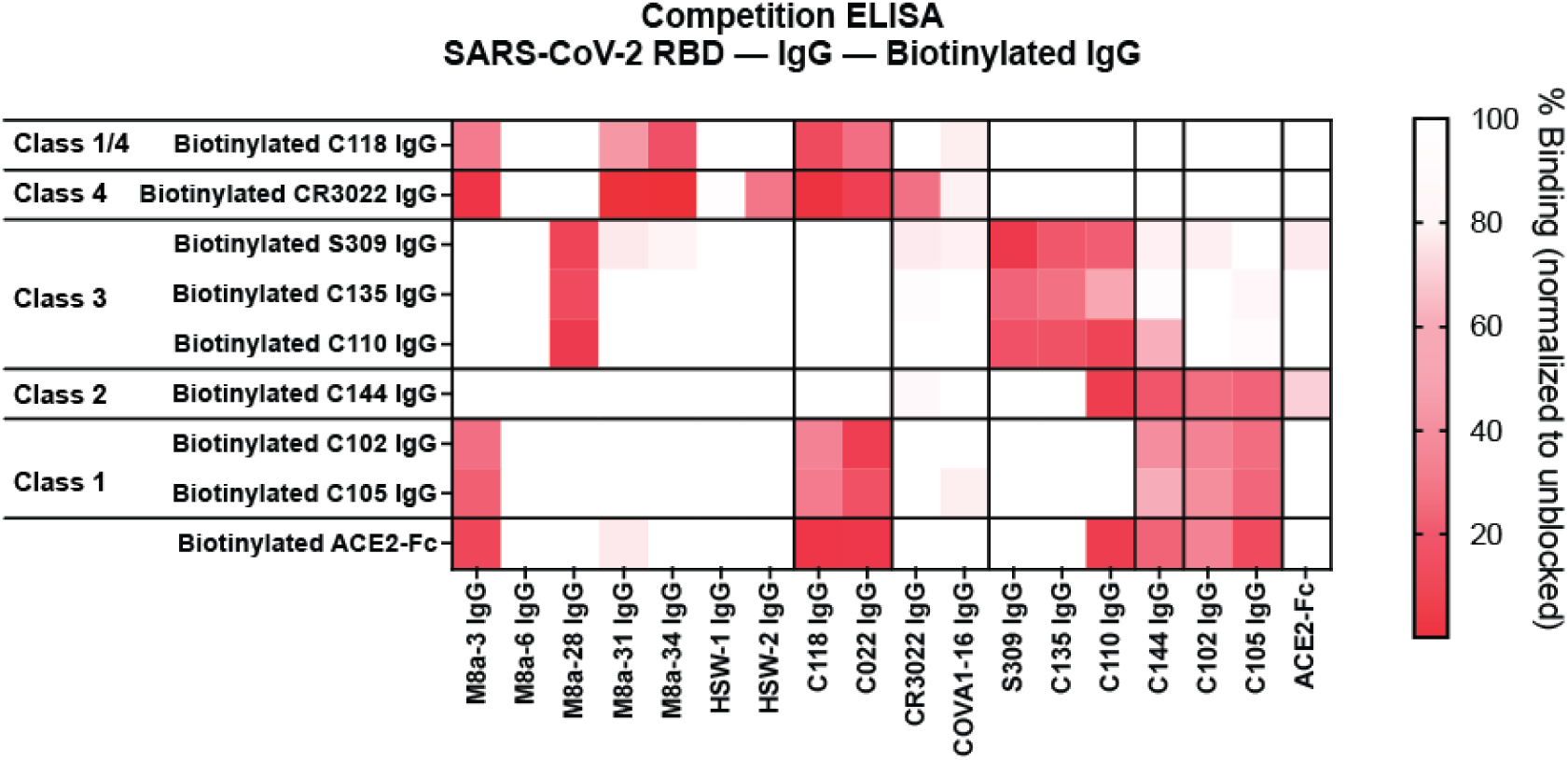
mAb epitope mapping. Competition ELISA experiment in which a test set of biotinylated IgGs of known epitopes or human ACE2-Fc (y-axis) were assayed for binding to SARS-CoV-2 RBD in the presence of unlabeled M8a, HSW, control IgGs, or ACE2-Fc (x-axis). The heat map shows shades of red indicating the percent binding of the tested IgGs or ACE2-Fc in the presence of competitor. M8a-3, M8a-31, M8a-34, and HSW-2 IgGs competed with biotinylated class 1/4 and/or class 4 anti-RBD, with HSW-2 competing with the biotinylated class 4 antibody only. M8a-3 IgG competed with biotinylated class 1 antibodies and ACE2-Fc in addition to the biotinylated class 1/4 and class 4 antibodies. M8a-28 IgG competed with biotinylated class 3 antibodies. M8a-6 and HSW-1 IgGs showed no competition in this assay.

**Figure S6.**
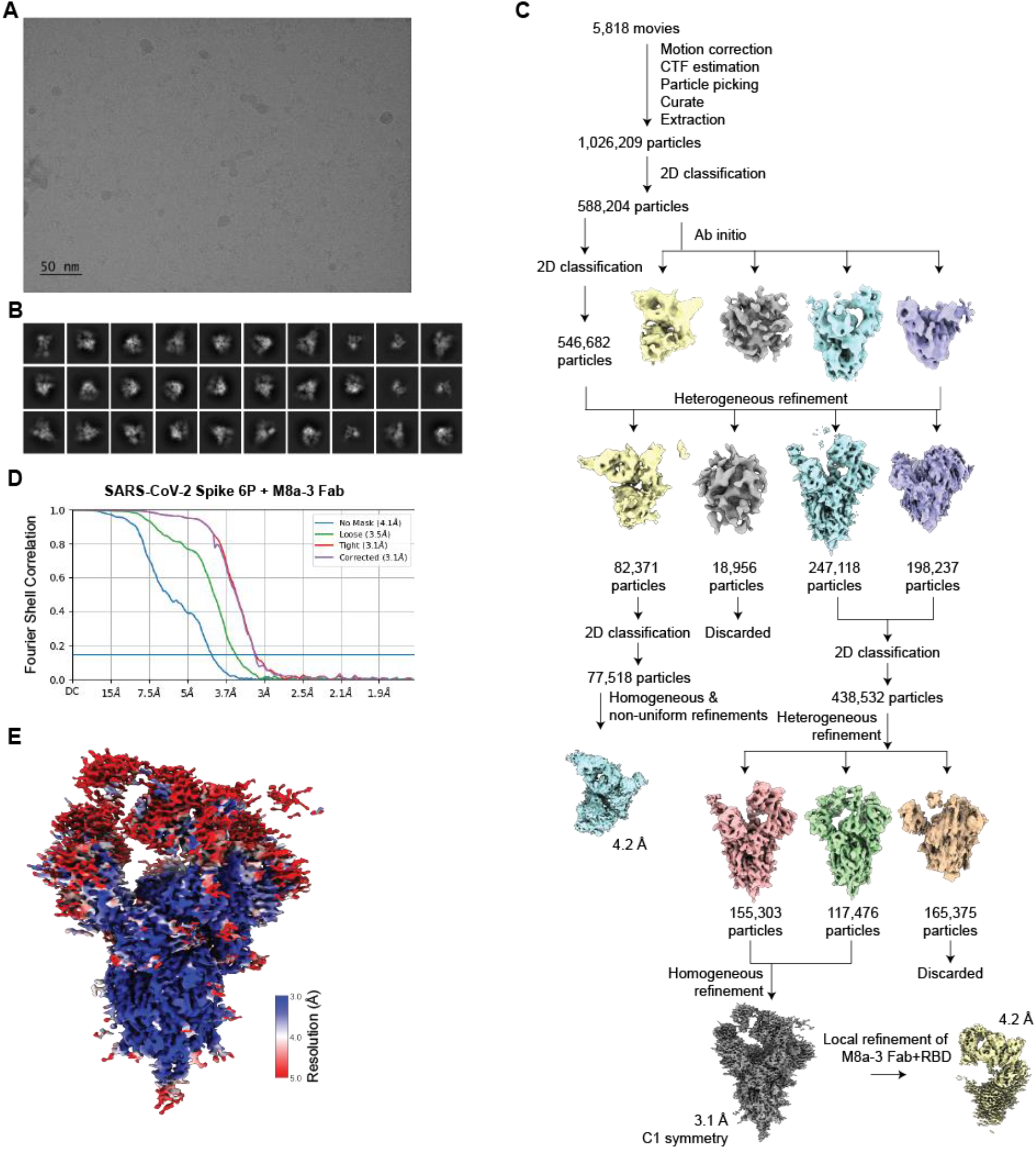
Cryo-EM data processing and validation for of M8a-3 Fab in complex with SARS-CoV-2 WA1 spike. (**A**) Representative micrograph. (**B**) Representative 2D classes. (**C**) Workflow of single-particle data processing. (**D**) Fourier shell correlation (FSC) plot of the final reconstruction. (**E**) Final reconstruction of M8a-3 Fab in complex with SARS-CoV-2 spike, colored by local resolution.

**Figure S7.**
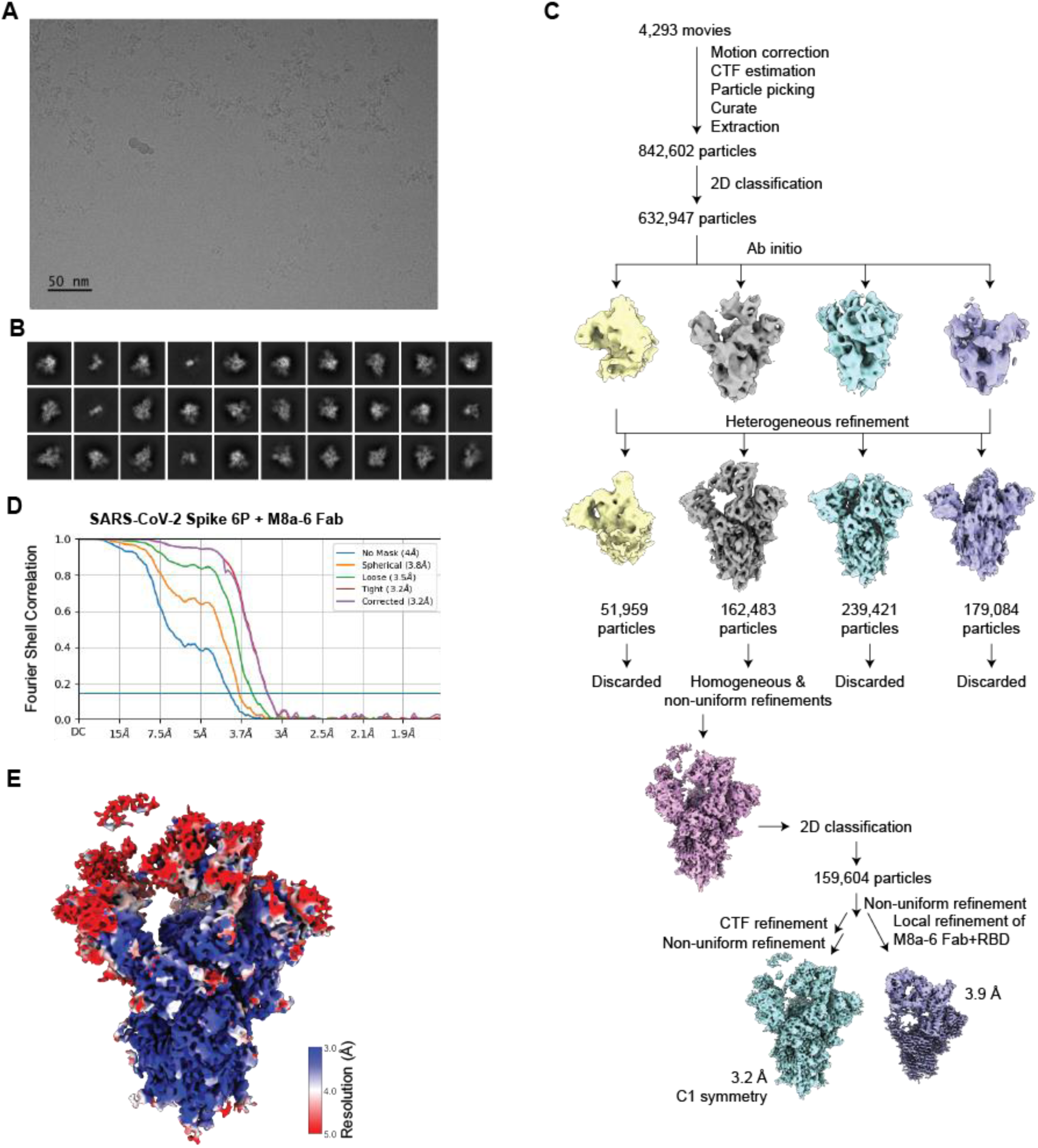
Cryo-EM data processing and validation for M8a-6 Fab in complex with SARS-CoV-2 WA1 spike. (**A**) Representative micrograph. (**B**) Representative 2D classes. (**C**) Workflow of single-particle data processing. (**D**) FSC plot of the final reconstruction. (**E**) Final reconstruction of M8a-6 Fab in complex with SARS-CoV-2 spike, colored by local resolution.

**Figure S8.**
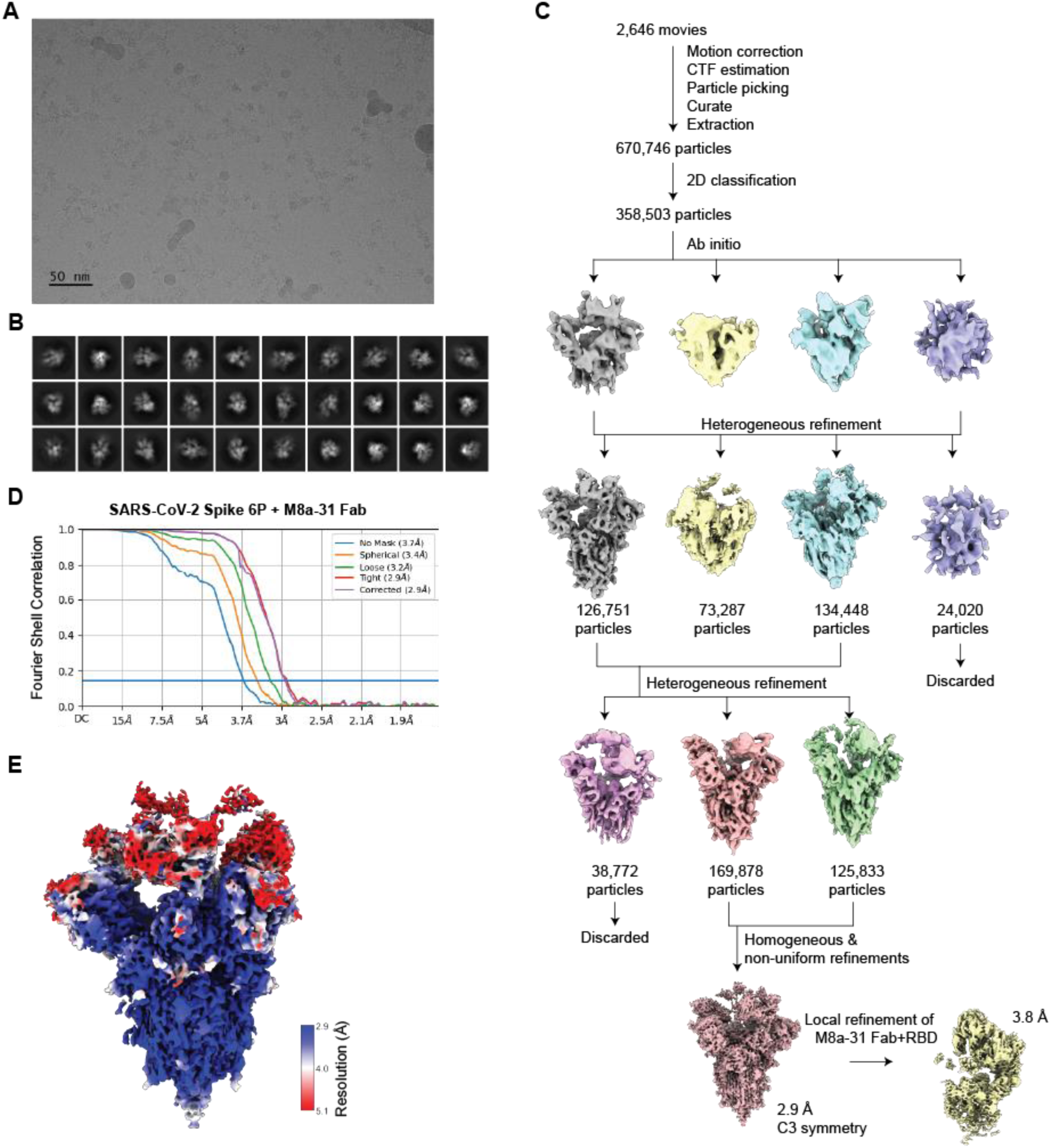
Cryo-EM data processing and validation for of M8a-31 Fab in complex with SARS-CoV-2 WA1 spike. (**A**) Representative micrograph. (**B**) Representative 2D classes. (**C**) Workflow of single-particle data processing. (**D**) FSC plot of the final reconstruction. (**E**) Final reconstruction of M8a-31 Fab in complex with SARS-CoV-2 spike, colored by local resolution.

**Figure S9.**
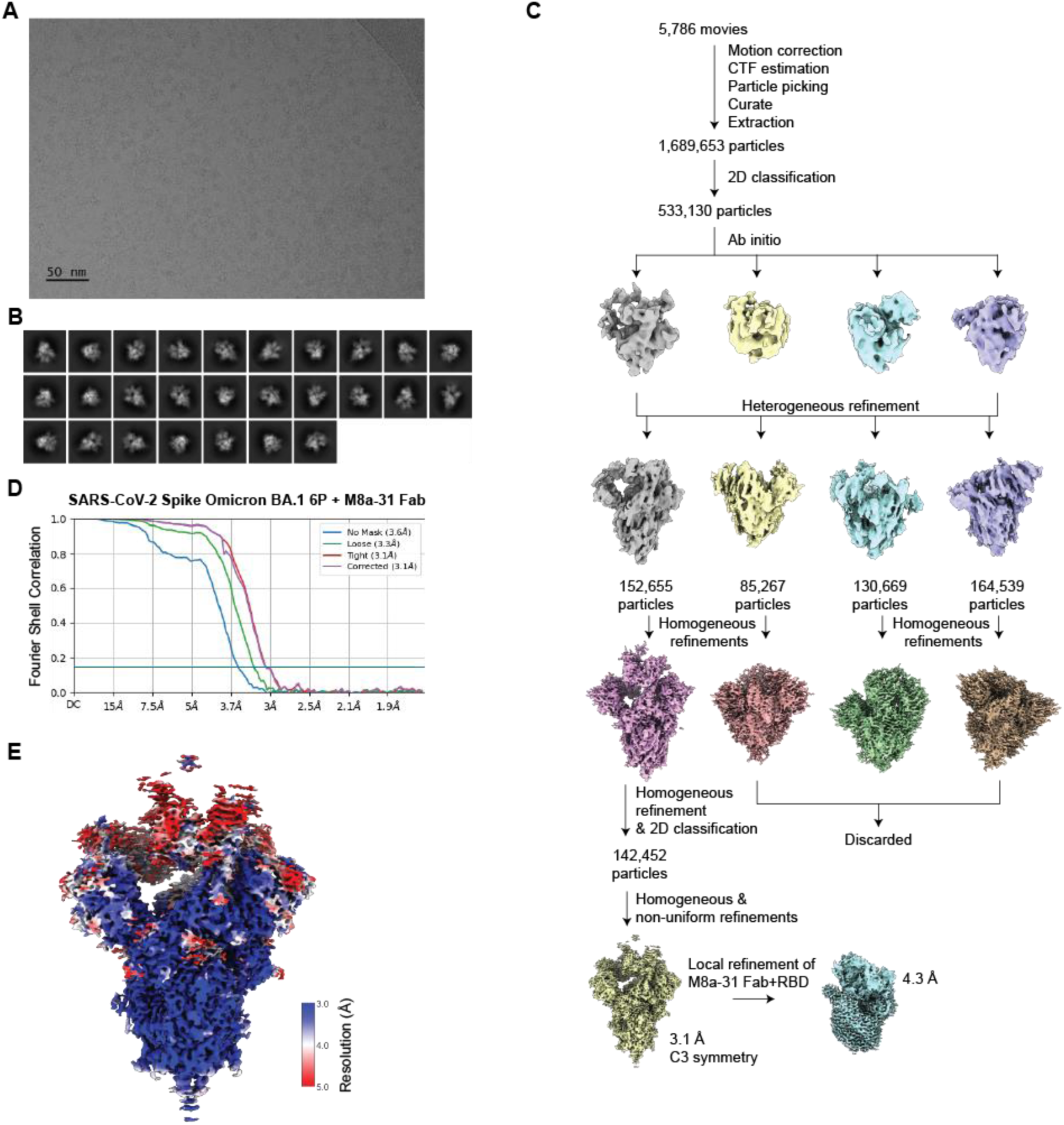
Cryo-EM data processing and validation for of M8a-31 Fab in complex with SARS-CoV-2 Omicron BA.1 spike. (**A**) Representative micrograph. (**B**) Representative 2D classes. (**C**) Workflow of single-particle data processing. (**D**) FSC plot of the final reconstruction. (**E**) Final reconstruction of M8a-3 Fab in complex with SARS-CoV-2 spike Omicron BA.1, colored by local resolution.

**Figure S10.**
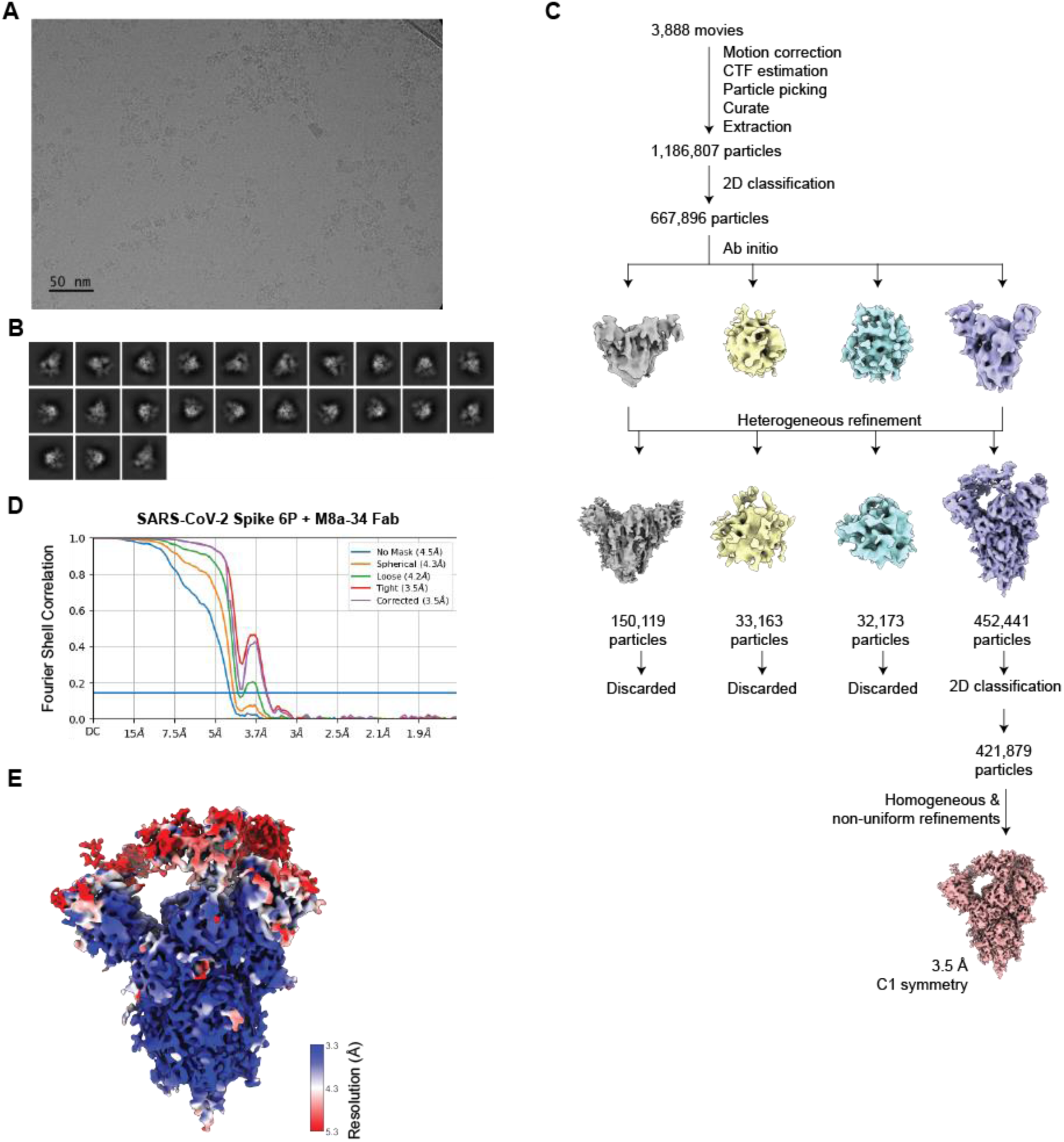
Cryo-EM data processing and validation for of M8a-34 Fab in complex with SARS-CoV-2 WA1 spike. (**A**) Representative micrograph. (**B**) Representative 2D classes. (**C**) Workflow of single-particle data processing. (**D**) FSC plot of the final reconstruction. (**E**) Final reconstruction of M8a-34 Fab in complex with SARS-CoV-2 spike, colored by local resolution.

**Figure S11.**
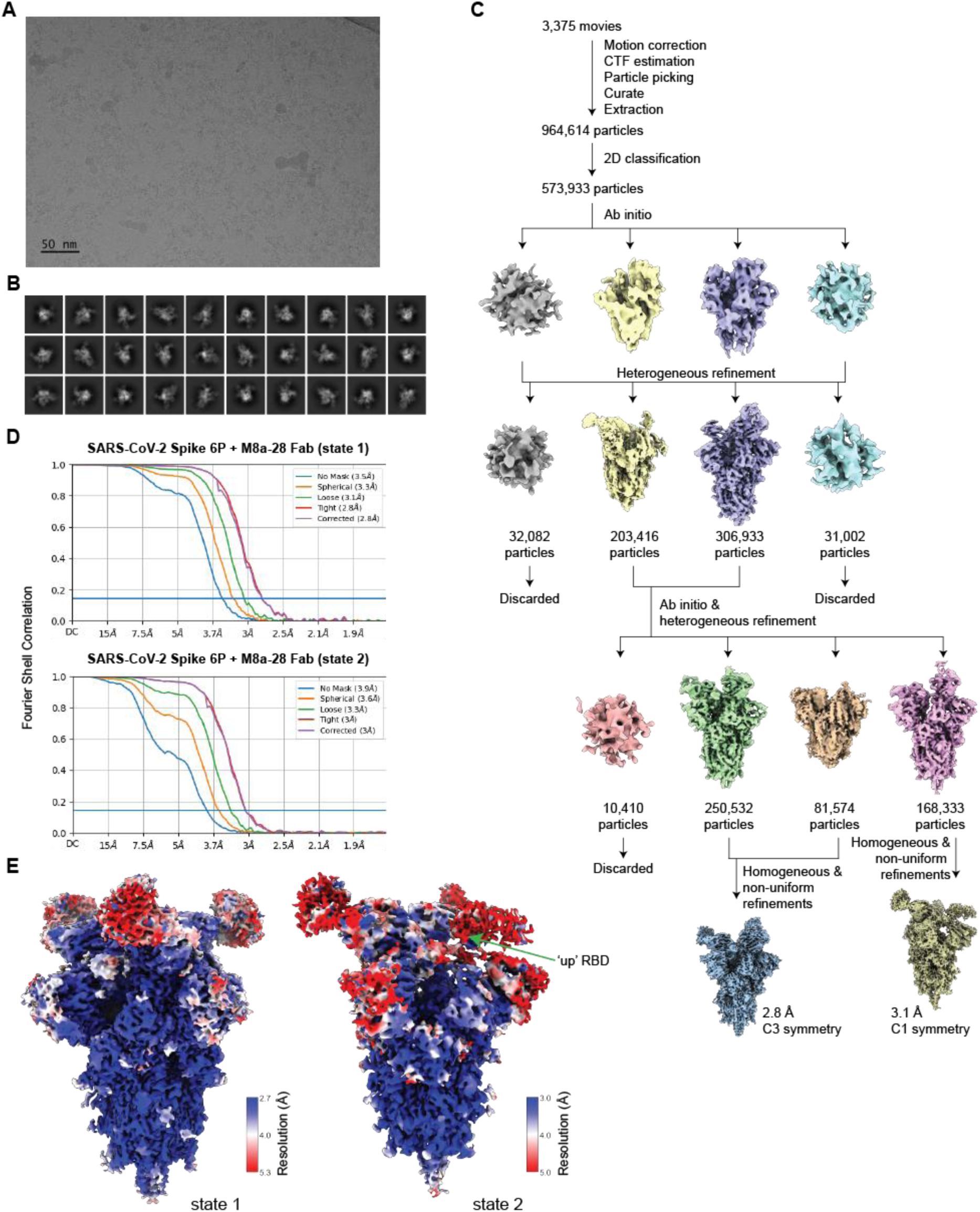
Cryo-EM data processing and validation for M8a-28 Fab in complex with SARS-CoV-2 WA1 spike. (**A**) Representative micrograph. (**B**) Representative 2D classes. (**C**) Workflow of single-particle data processing. (**D**) FSC plot of the final reconstruction. (**E**) Final reconstructions of both states of M8a-28 Fab in complex with SARS-CoV-2 spike, colored by local resolution. An ‘up’ RBD in state 2 is labeled with an arrow.

**Figure S12.**
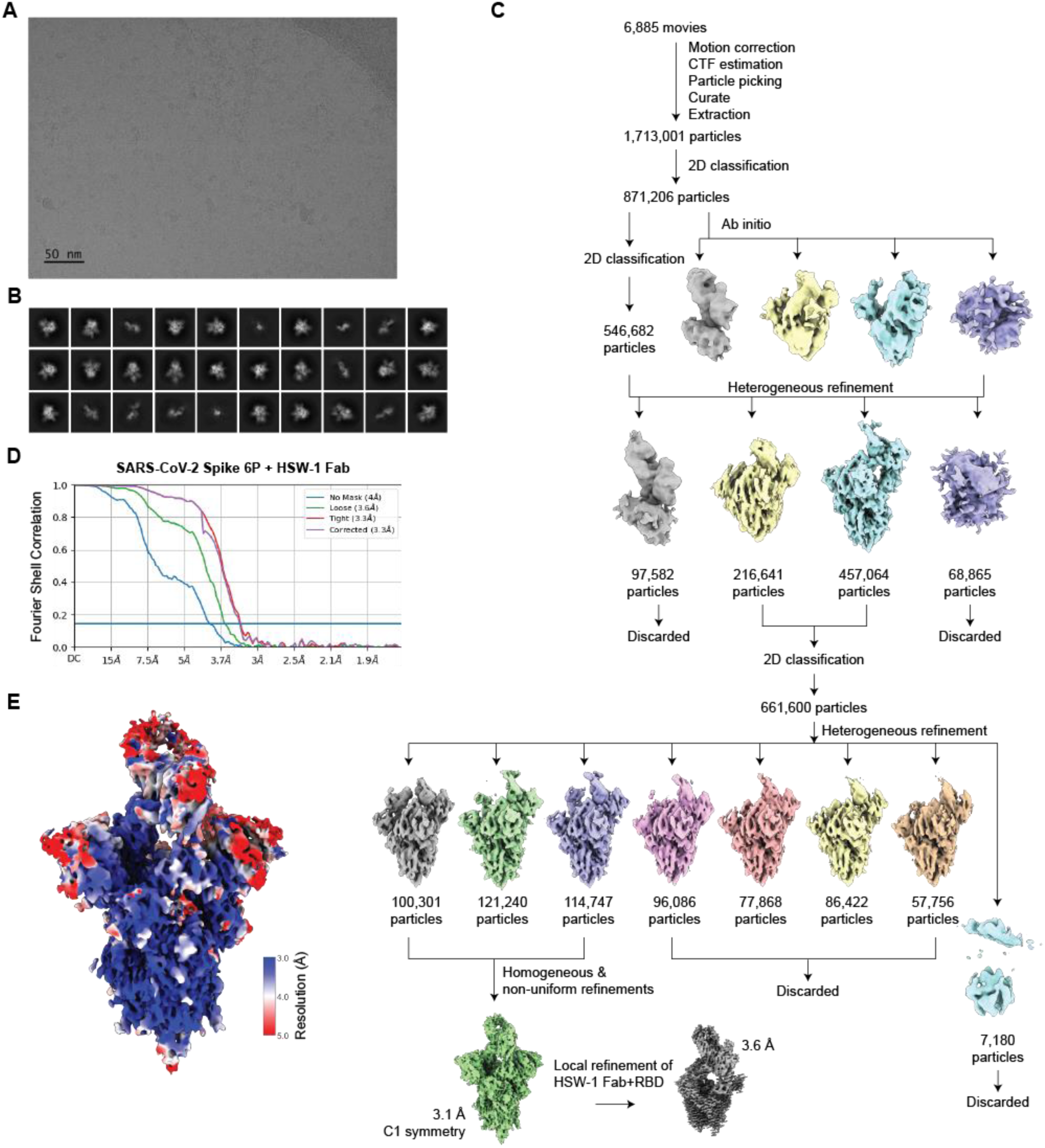
Cryo-EM data processing and validation for of HSW-1 Fab in complex with SARS-CoV-2 WA1 spike. (**A**) Representative micrograph. (**B**) Representative 2D classes. (**C**) Workflow of single-particle data processing. (**D**) FSC plot of the final reconstruction. (**E**) Final reconstruction of HSW-1 Fab in complex with SARS-CoV-2 spike, color by local resolution.

**Figure S13.**
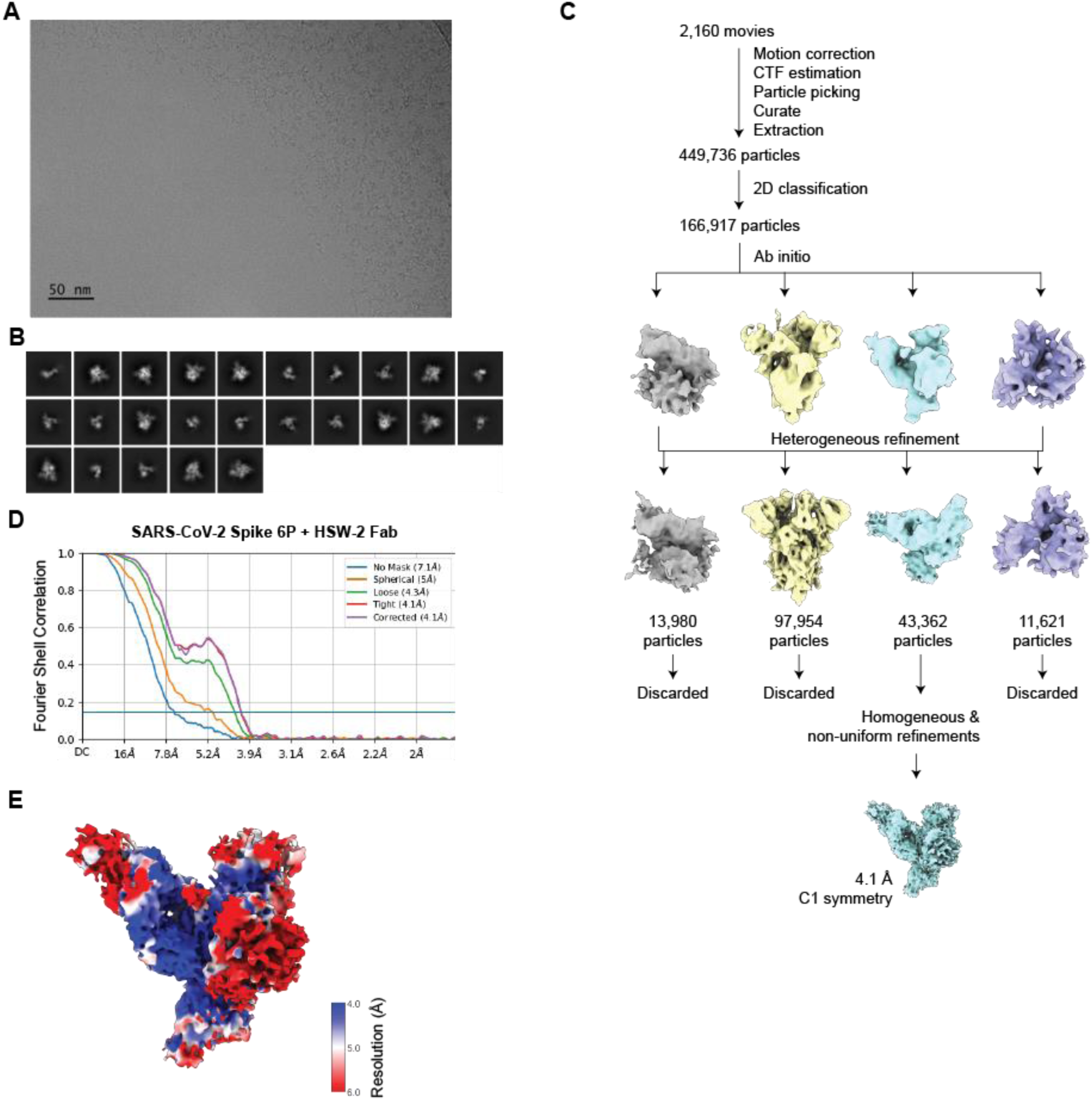
Cryo-EM data processing and validation for of HSW-2 Fab in complex with SARS-CoV-2 WA1 spike. (**A**) Representative micrograph. (**B**) Representative 2D classes. (**C**) Workflow of single-particle data processing. (**D**) FSC plot of the final reconstruction. (**E**) Final reconstruction of HSW-2 Fab in complex with SARS-CoV-2 spike S1 domain, color by local resolution.

**Figure S14.**
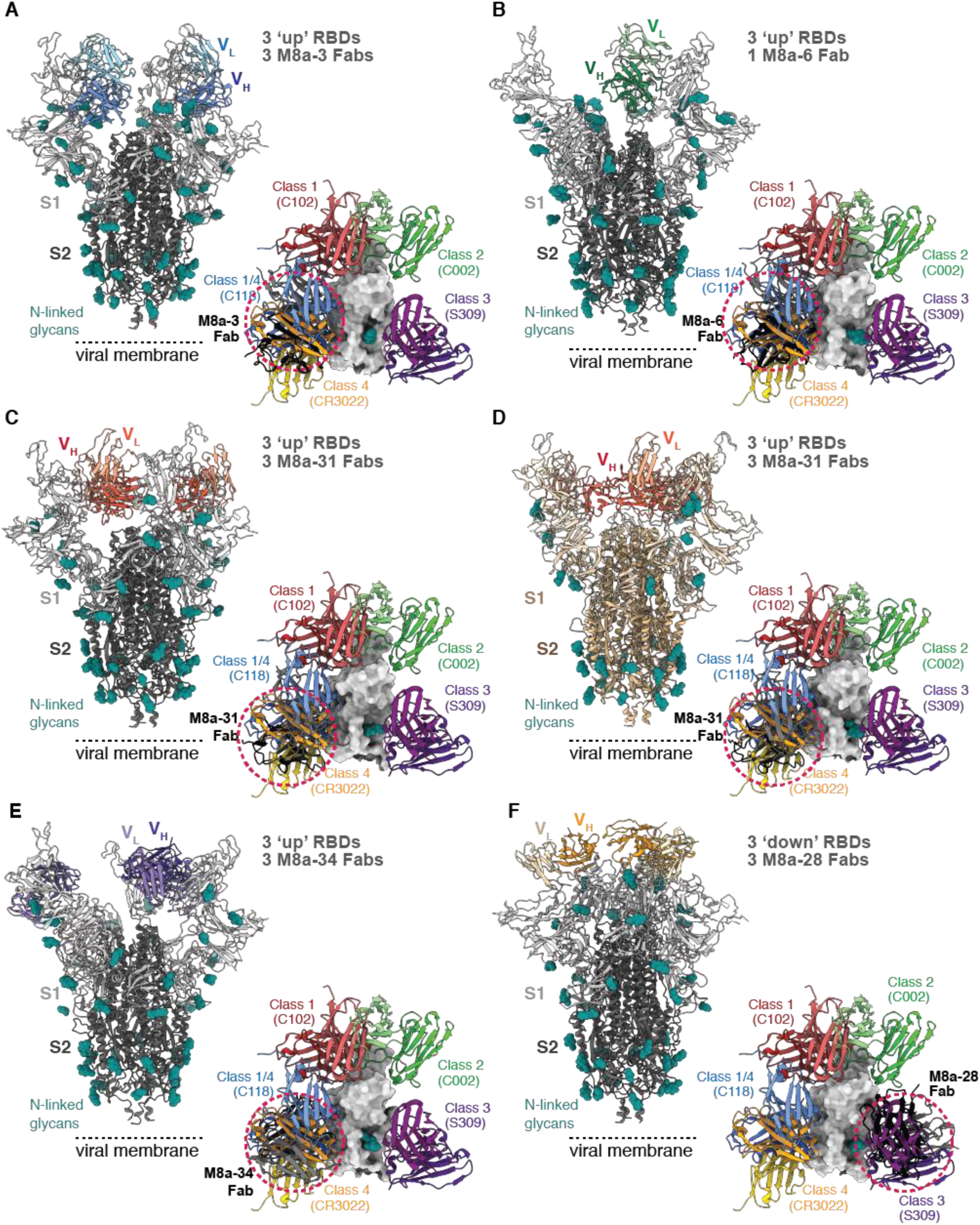
mAbs isolated from mice immunized with mosaic-8 nanoparticles. Cartoon representations of single particle cryo-EM structures of Fab-spike trimer complexes are shown from the side (left) with a comparison of binding epitopes of the Fab with representative anti-RBD antibodies: class 1 (C102, PDB 7K8M), class 2 (C002, PDB 7K8T), class 3 (S309, PDB 7JX3), class 4 (CR3022, PDB 7LOP), and class 1/4 (C118, PDB 7RKV)) aligned on an RBD in surface representation (right). Only V_H_-V_L_ domains are shown for each Fab. Fabs of interests (colored in black and circled with a red dotted line) and the anti-RBD antibodies used for classification are aligned on a surface representation of the RBD. N-linked glycans are shown as teal spheres. (**A**) WA1 spike complexed with M8a-3. (**B**) SARS-CoV-2 WA1 spike complexed with M8a-6. (**C**) WA1 spike complexed with M8a-31. (**D**) Omicron BA.1 spike complexed with M8a-31. (**E**) WA1 spike complexed with M8a-34. (**F**) WA1 spike complexed with M8a-28.

**Figure S15.**
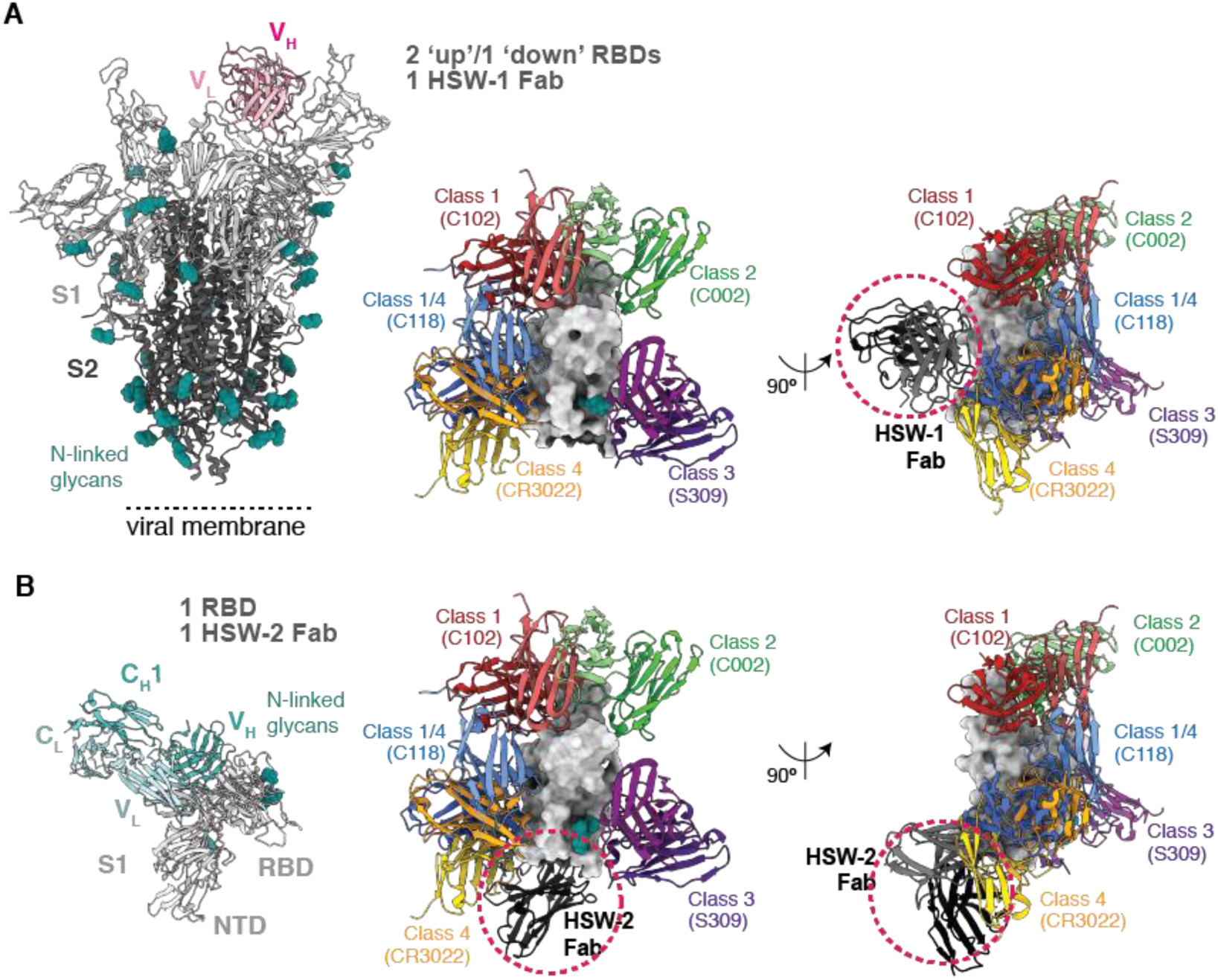
mAbs isolated from mice immunized with homotypic nanoparticles. Cartoon representations of single particle cryo-EM structures of Fab-spike trimer complexes are shown from the side (left) with a comparison of binding epitopes of the Fab with representative anti-RBD antibodies: class 1 (C102, PDB 7K8M), class 2 (C002, PDB 7K8T), class 3 (S309, PDB 7JX3), class 4 (CR3022, PDB 7LOP) and class 1/4 (C118, PDB 7RKV)) aligned on an RBD in surface representation (middle and right). Only V_H_-V_L_ domains are shown for each Fab. Fabs of interests (colored in black and circled with a red dotted line) and the anti-RBD antibodies used for classification are aligned on a surface representation of the RBD. N-linked glycans are shown as teal spheres. (**A**) SARS-CoV-2 WA1 spike complexed with HSW-1. (**B**) WA1 spike S1 domain complexed with HSW-2.

**Figure S16.**
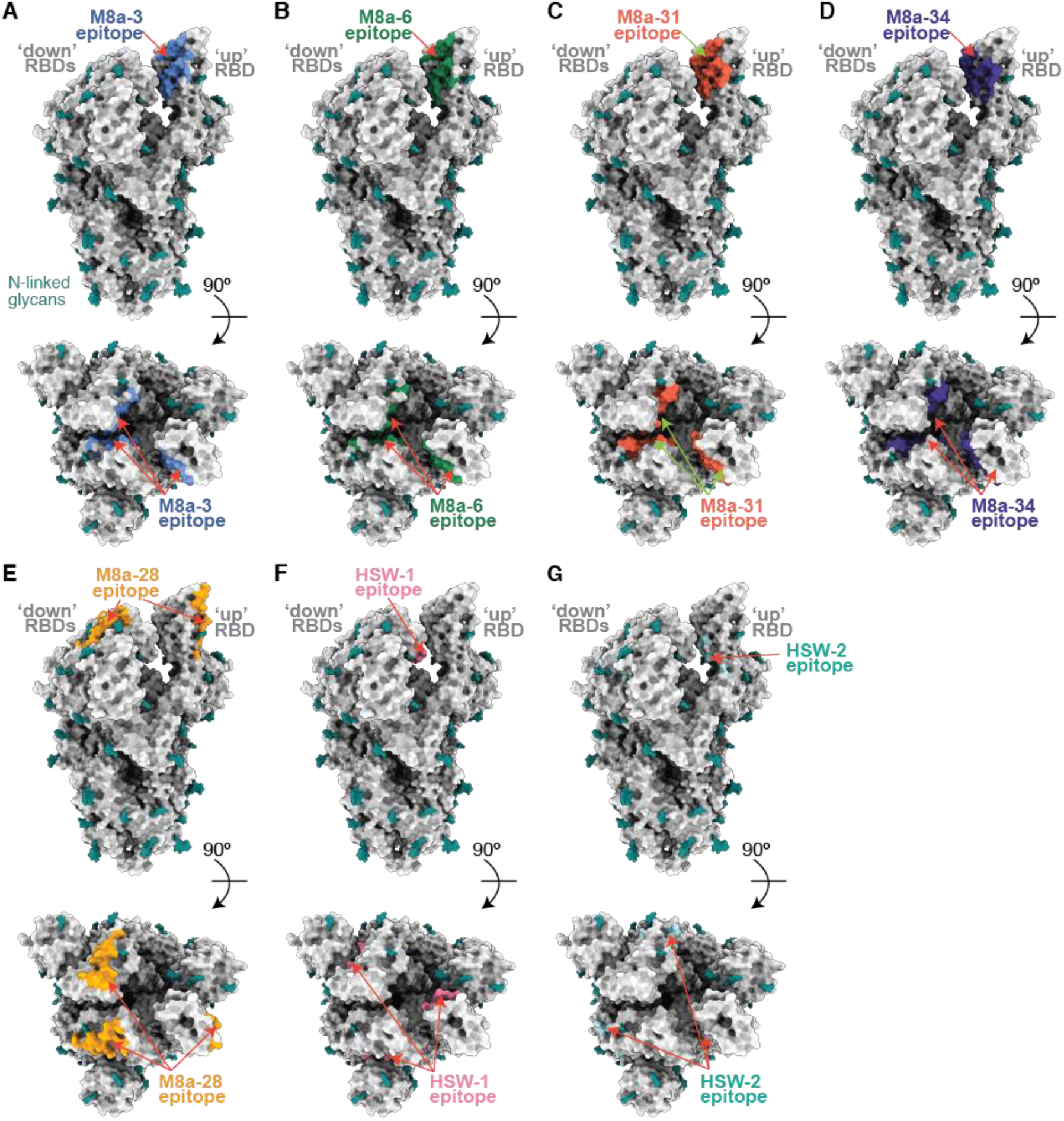
Epitopes of mAbs mapped on a unliganded SARS-CoV-2 spike trimer. Binding epitopes for all mAbs were identified by PDBePISA (Krissinel and Henrick, 2007) and then mapped on to a unliganded spike trimer with two ‘down’ and one ‘up’ RBDs (PDB 6VYB). The spike trimer is shown as a surface representation with N-linked glycans shown as teal spheres. The epitopes of (**A**) M8a-3, (**B**) M8a-6, (**C**) M8a-31 and (**D**) M8a-34 are blocked in the ‘down’ RBD conformation, but accessible in an ‘up’ RBD conformation. (**E**) The epitope of M8a-28 is accessible in both ‘down’ and ‘up’ RBD conformations. (**F**) The epitope of HSW-1 is blocked in the ‘down’ RBD conformation, but accessible in an ‘up’ RBD conformation. (**G**) The epitope of HSW-2 is blocked in both ‘down’ and ‘up’ RBD conformations.

**Figure S17.**
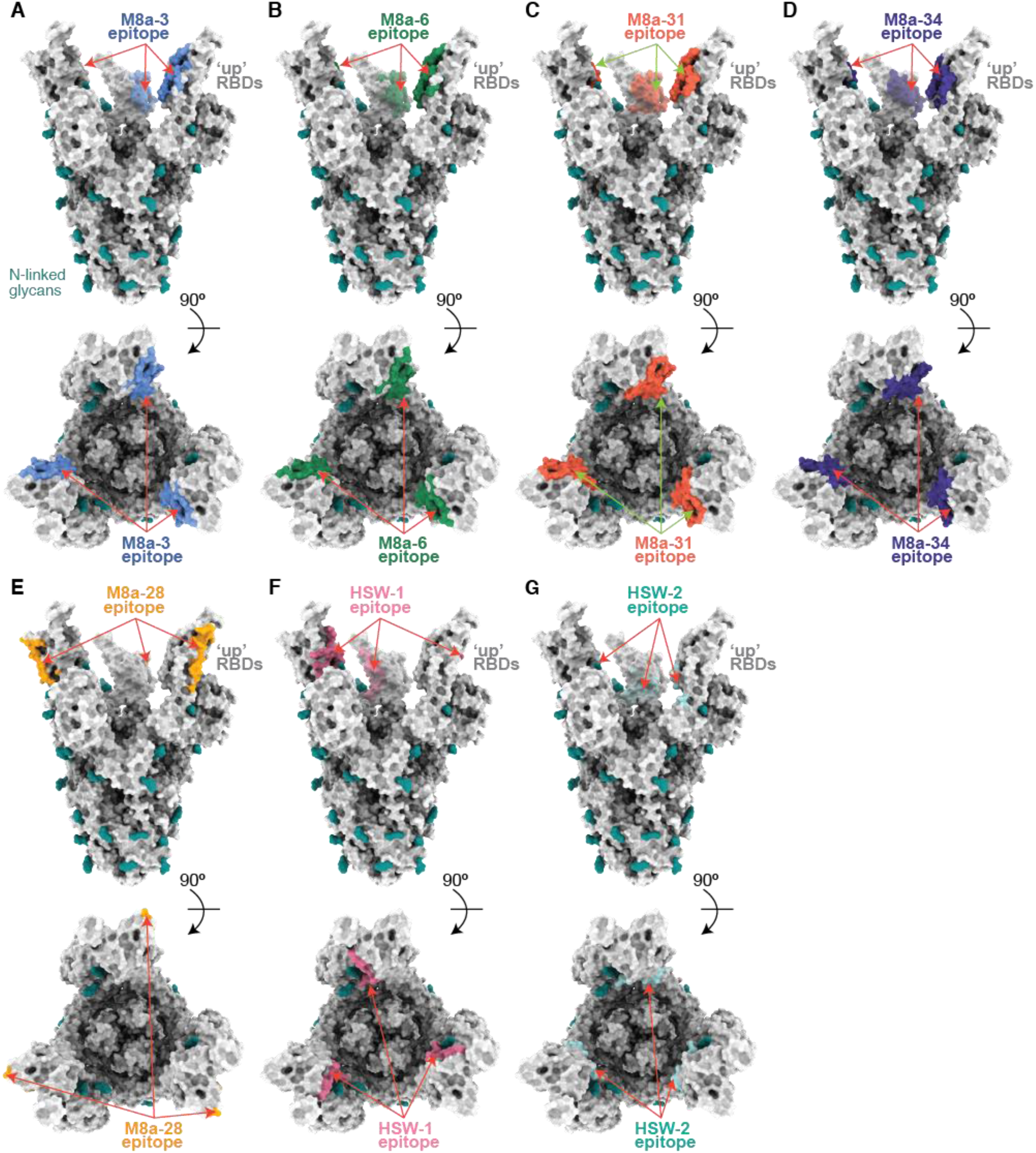
Epitopes of mAbs mapped on a SARS-CoV-2 spike trimer with all ‘up’ RBDs. Binding epitopes for all mAbs were identified by PDBePISA (Krissinel and Henrick, 2007) and then mapped on to a spike trimer with three ‘up’ RBDs (PDB 7RKV; Fabs not shown). The spike trimer is shown as a surface representation with N-linked glycans shown as teal spheres. The epitopes of (**A**) M8a-3, (**B**) M8a-6, (**C**) M8a-31, and (**D**) M8a-34 are all accessible in an ‘up’ RBD conformation. (**E**) The epitope of M8a-28 is accessible in ‘up’ RBD conformation. (**F**) The epitope of HSW-1 is accessible in an ‘up’ RBD conformation. (**G**) The epitope of HSW-2 is sterically hindered in an ‘up’ RBD conformation.

**Figure S18.**
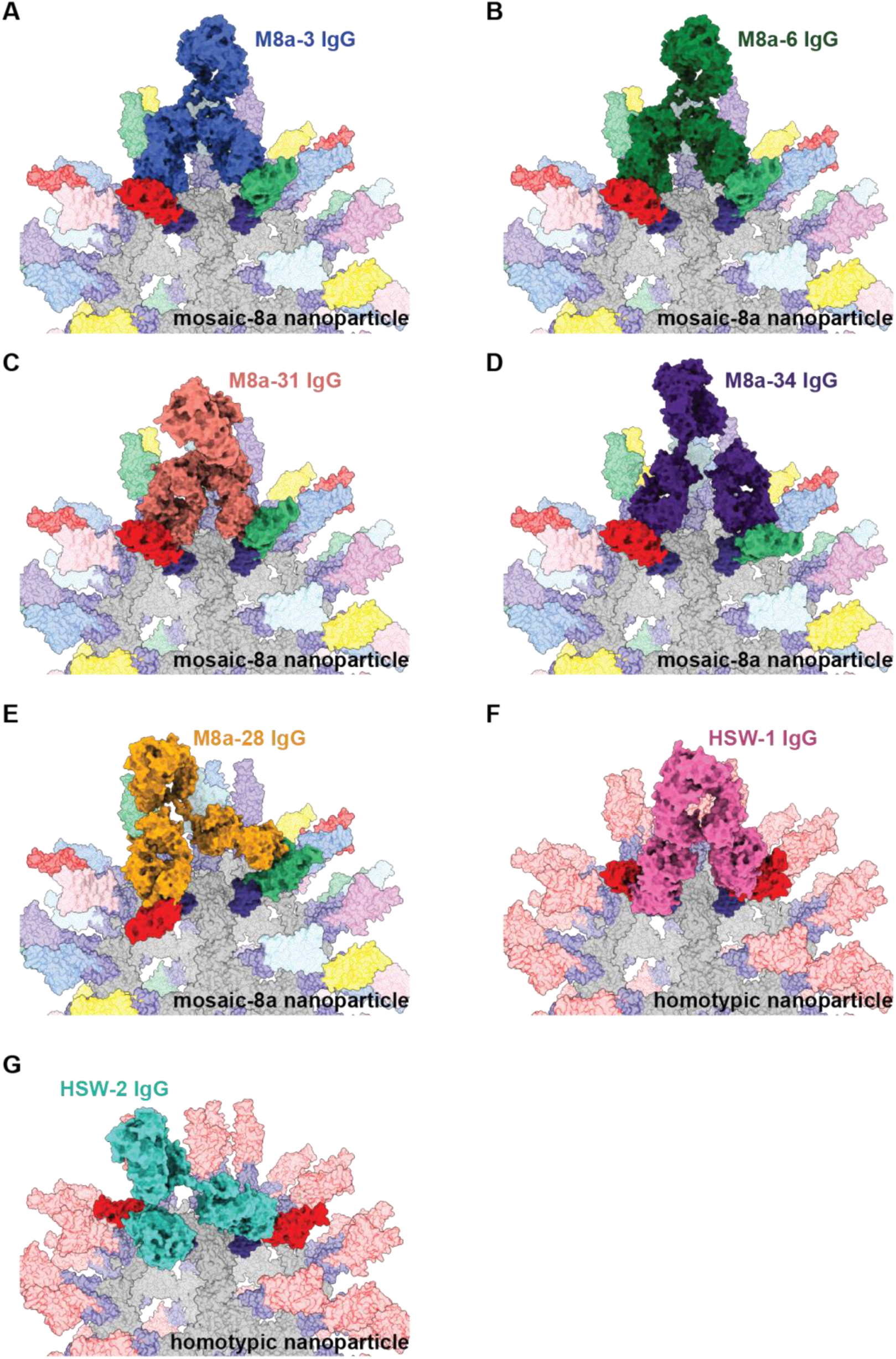
Models of M8a and HSW IgGs binding to adjacent RBDs on SpyCatcher-mi3 nanoparticles. Structural models of RBD-nanoparticles formed by SpyCatcher-mi3 and SpyTagged RBDs were made using coordinates of an RBD (PDB 7SC1) (represented in different colors for mosaic-8 nanoparticles, with two adjacent RBDs in red and blue, or in red for homotypic nanoparticles), mi3 (PDB 7B3Y) (gray), and SpyCatcher (PDB 4MLI) (dark blue). IgGs (various colors) were modeled using the coordinates of each mAb Fab based on an intact IgG crystal structure (PDB 1HZH), and the RBD binding epitope of each Fab in a modeled IgGs was determined based on the Fab-spike structures reported in this study. The two Fabs of each IgGs were positioned with distances between the C-termini of the Fab C_H_1 domains to be less than 65 Å, as described previously (Barnes et al., 2020a). Both the IgG Fc hinge and the linker region between a SpyTagged RBD and SpyCatcher were assumed to be flexible and adjusted accordingly. Models are shown for (**A**) M8a-3 IgG interacting with two adjacent RBDs on a mosaic-8 RBD-nanoparticle. (**B**) M8a-6 IgG interacting with two adjacent RBDs on a mosaic-8 RBD-nanoparticle. (**C**) M8a-31 IgG interacting with two adjacent RBDs on a mosaic-8 RBD-nanoparticle. (**D**) M8a-34 IgG interacting with two adjacent RBDs on a mosaic-8 RBD-nanoparticle. (**E**) M8a-28 IgG interacting with two adjacent RBDs on a mosaic-8 RBD-nanoparticle. (**F**) HSW-1 IgG interacting with two adjacent RBDs on a homotypic RBD-nanoparticle. (**G**) HSW-2 IgG interacting with two adjacent RBDs on a homotypic RBD-nanoparticle.

**Table S1.**
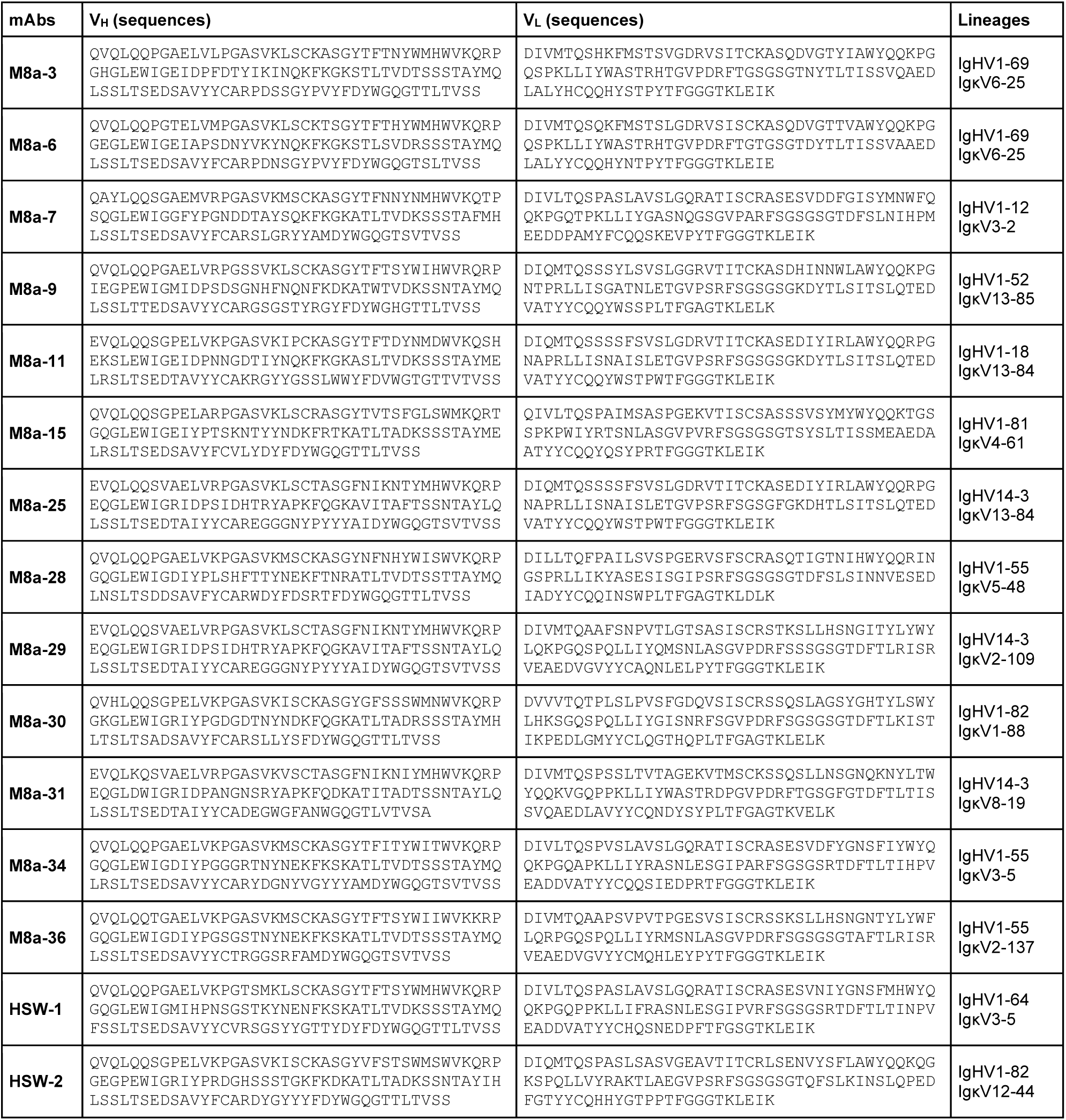
Sequences and V gene segment lineages for 16 mAbs identified as binding at least one RBD during screening. M8a-9 and M8a-36 did not exhibit binding to purified RBDs by ELISA (data not shown). M8a-11 and M8a-26 are identical sequences.

**Table S2.** Single-particle cryo-EM data collection, processing, and refinement.

| Structures | WA1 spike in complex with |  |  |  |  |  |  | Omicron |
| --- | --- | --- | --- | --- | --- | --- | --- | --- |
|  | M8a-3 Fab | M8a-6 Fab | M8a-28 Fab | M8a-31 Fab | M8a-34 Fab | HSW-1 Fab | HSW-2 Fab | BA.1 spike in complex with M8a-31 Fab |
| Data Collection and processing |  |  |  |  |  |  |  |  |
| Microscope | Titan Krios | Titan Krios | Titan Krios | Titan Krios | Titan Krios | Titan Krios | Talos Arctica | Titan Krios |
| Camera | Gatan K3 | Gatan K3 | Gatan K3 | Gatan K3 | Gatan K3 | Gatan K3 | Gatan K3 | Gatan K3 |
| Magnification | 105,000 | 105,000 | 105,000 | 105,000 | 105,000 | 105,000 | 45,000 | 105,000 |
| Voltage (keV) | 300 | 300 | 300 | 300 | 300 | 300 | 200 | 300 |
| Exposure (e/Å²) | 60 | 60 | 60 | 60 | 60 | 60 | 60 | 60 |
| Pixel size (Å) | 0.832 | 0.832 | 0.832 | 0.832 | 0.832 | 0.832 | 0.869 | 0.832 |
| Defocus Range (µm) | -1 to -3 | -1 to -3 | -1 to -3 | -1 to -3 | -1 to -3 | -1 to -3 | -1 to -3 | -1 to -3 |
| Initial Particle Image (no.) | 1,026,209 | 842,602 | 964,614 | 670,746 | 1,186,807 | 1,713,001 | 449,736 | 1,689,653 |
| Final Particle Image (no.) | 272,779 | 159,604 | 332,106 | 295,711 | 421,879 | 336,288 | 43,362 | 142,452 |
| Symmetry Imposed | C1 | C1 | C3 | C3 | C1 | C1 | C1 | C3 |
| Map Resolution (Å) | 3.1 | 3.2 | 2.8 | 2.9 | 3.5 | 3.1 | 4.1 | 3.1 |
| FSC Threshold | 0.143 | 0.143 | 0.143 | 0.143 | 0.143 | 0.143 | 0.143 | 0.143 |
| Map Resolution Range (Å) | 3.0 - 3.5 | 3.1 - 3.5 | 2.7 to 3.1 | 2.9 - 3.2 | 3.5 - 4.3 | 3.0 to 3.4 | 3.9 - 4.3 | 2.9 - 3.3 |
| Refinement |  |  |  |  |  |  |  |  |
| Initial Model Used | PDB 7SC1 | PDB 7SC1 | PDB 7SC1 | PDB 7SC1 | PDB 7SC1 | PDB 7SC1 | PDB 7SC1 | PDB 7SC1 |
| Model Resolution (Å) | 3.1 | 3.2 | 2.8 | 2.9 | 3.5 | 3.1 | 4.1 | 3.1 |
| FSC Threshold | 0.143 | 0.143 | 0.143 | 0.143 | 0.143 | 0.143 | 0.143 | 0.143 |
| Model composition |  |  |  |  |  |  |  |  |
| non-hydrogen atoms | 30,762 | 27,051 | 30,942 | 30,552 | 30,669 | 26,992 | 8,392 | 30,075 |
| protein residues | 3,867 | 3,397 | 3,906 | 3,861 | 3,873 | 3,409 | 1,077 | 3,801 |
| ligands | 51 | 48 | 39 | 45 | 42 | 38 | 4 | 33 |
| Average B-factors (Å²) |  |  |  |  |  |  |  |  |
| protein | 135.2 | 157.5 | 110.4 | 130.4 | 205.7 | 158.8 | 197.9 | 92.4 |
| ligands | 116.3 | 139.9 | 96.8 | 111.6 | 167.3 | 143.6 | 213.5 | 82.5 |
| R.m.s. deviations |  |  |  |  |  |  |  |  |
| Bond length (Å) | 0.005 | 0.004 | 0.003 | 0.004 | 0.003 | 0.004 | 0.003 | 0.004 |
| Bond angles (°) | 0.562 | 0.582 | 0.536 | 0.572 | 0.586 | 0.593 | 0.589 | 0.594 |
| Validation |  |  |  |  |  |  |  |  |
| MolProbity score | 1.85 | 1.63 | 1.74 | 1.72 | 1.75 | 1.83 | 1.9 | 1.77 |
| Clashscore | 10.3 | 9.4 | 9.5 | 9.1 | 11.1 | 12.3 | 14.6 | 10.8 |
| Rotamer outliers | 0.06 | 0.07 | 0.03 | 0 | 0.03 | 0 | 0 | 0.06 |
| Ramachandran plot |  |  |  |  |  |  |  |  |
| Ramachandran favored (%) | 95.4 | 97.3 | 96.4 | 96.5 | 96.9 | 96.5 | 96.4 | 96.6 |
| Ramachandran allowed (%) | 4.6 | 2.7 | 3.6 | 3.5 | 3.1 | 3.5 | 3.6 | 3.4 |
| Ramachandran outliers (%) | 0 | 0 | 0 | 0 | 0 | 0 | 0 | 0 |
| PDB ID | 7UZ4 | 7UZ5 | 7UZ6 | 7UZ7 | 7UZ9 | 7UZA | 7UZB | 7UZ8 |

**Table S3.**
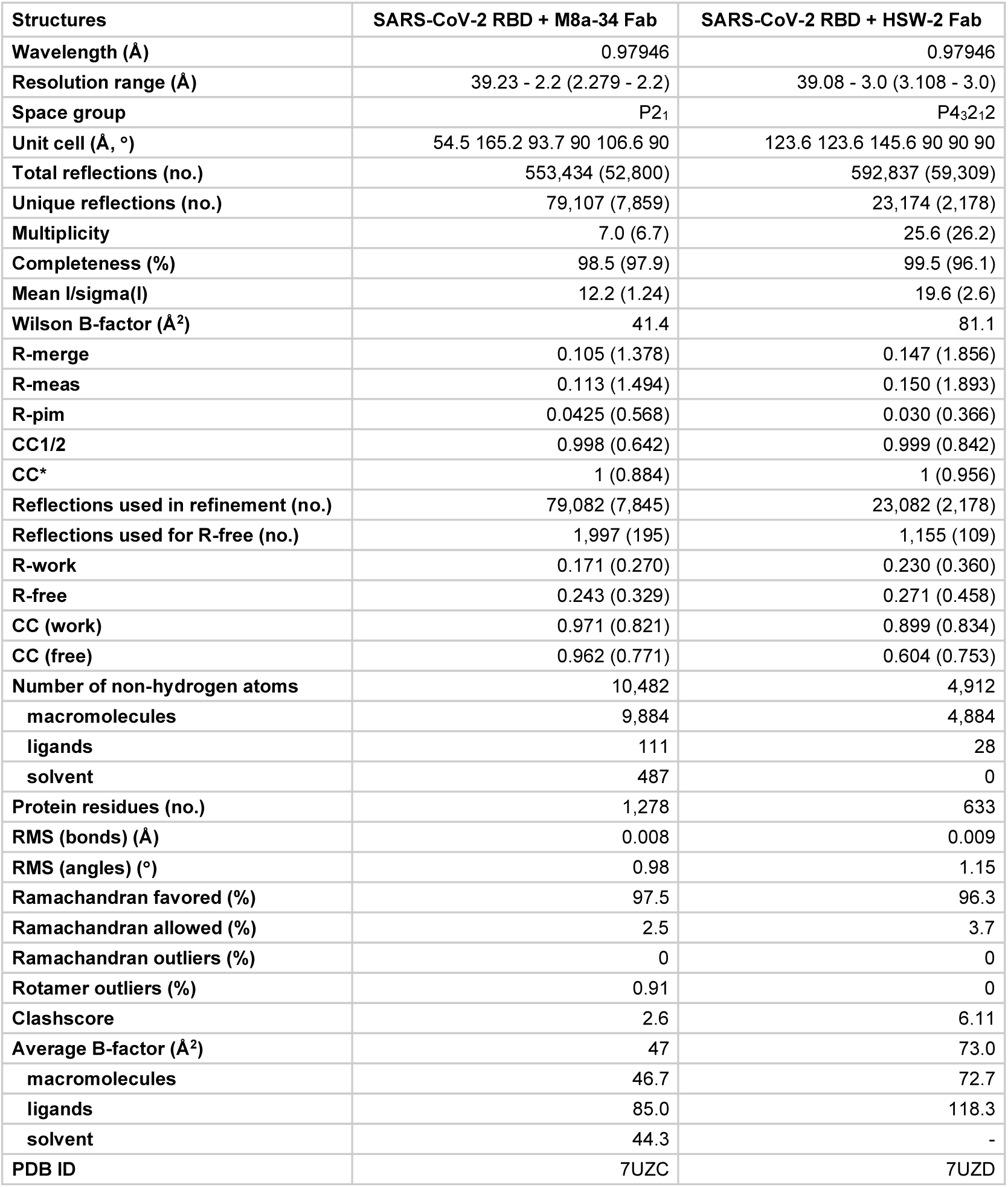
X-ray crystallography data collection, processing, and refinement.

| Structures | SARS-CoV-2 RBD + M8a-34 Fab | SARS-CoV-2 RBD + HSW-2 Fab |
| --- | --- | --- |
| Wavelength (Å) | 0.97946 | 0.97946 |
| Resolution range (Å) | 39.23 - 2.2 (2.279 - 2.2) | 39.08 - 3.0 (3.108 - 3.0) |
| Space group | P2<sub>1</sub> | P4<sub>3</sub>2<sub>1</sub>2 |
| Unit cell (Å, °) | 54.5 165.2 93.7 90 106.6 90 | 123.6 123.6 145.6 90 90 90 |
| Total reflections (no.) | 553,434 (52,800) | 592,837 (59,309) |
| Unique reflections (no.) | 79,107 (7,859) | 23,174 (2,178) |
| Multiplicity | 7.0 (6.7) | 25.6 (26.2) |
| Completeness (%) | 98.5 (97.9) | 99.5 (96.1) |
| Mean I/sigma(I) | 12.2 (1.24) | 19.6 (2.6) |
| Wilson B-factor (Å<sup>2</sup>) | 41.4 | 81.1 |
| R-merge | 0.105 (1.378) | 0.147 (1.856) |
| R-meas | 0.113 (1.494) | 0.150 (1.893) |
| R-pim | 0.0425 (0.568) | 0.030 (0.366) |
| CC1/2 | 0.998 (0.642) | 0.999 (0.842) |
| CC* | 1 (0.884) | 1 (0.956) |
| Reflections used in refinement (no.) | 79,082 (7,845) | 23,082 (2,178) |
| Reflections used for R-free (no.) | 1,997 (195) | 1,155 (109) |
| R-work | 0.171 (0.270) | 0.230 (0.360) |
| R-free | 0.243 (0.329) | 0.271 (0.458) |
| CC (work) | 0.971 (0.821) | 0.899 (0.834) |
| CC (free) | 0.962 (0.771) | 0.604 (0.753) |
| Number of non-hydrogen atoms | 10,482 | 4,912 |
| macromolecules | 9,884 | 4,884 |
| ligands | 111 | 28 |
| solvent | 487 | 0 |
| Protein residues (no.) | 1,278 | 633 |
| RMS (bonds) (Å) | 0.008 | 0.009 |
| RMS (angles) (°) | 0.98 | 1.15 |
| Ramachandran favored (%) | 97.5 | 96.3 |
| Ramachandran allowed (%) | 2.5 | 3.7 |
| Ramachandran outliers (%) | 0 | 0 |
| Rotamer outliers (%) | 0.91 | 0 |
| Clashscore | 2.6 | 6.11 |
| Average B-factor (Å<sup>2</sup>) | 47 | 73.0 |
| macromolecules | 46.7 | 72.7 |
| ligands | 85.0 | 118.3 |
| solvent | 44.3 | - |
| PDB ID | 7UZC | 7UZD |

